# Ribosome cooperativity promotes fast and efficient translation

**DOI:** 10.1101/2024.04.08.588515

**Authors:** Maximilian F. Madern, Sora Yang, Marvin E. Tanenbaum

## Abstract

The genetic information stored in mRNA is decoded by ribosomes during mRNA translation. mRNAs are typically translated by multiple ribosomes simultaneously, but it is unclear if different ribosomes translating the same mRNA functionally interact with each other. Here, we combine single-ribosome imaging and simulations to compare translation dynamics of individual ribosomes in either monosomes or polysomes to determine how ribosomes interact during translation. We find that ribosomes frequently undergo transient collisions, but that transient collisions, unlike persistent collisions, escape detection by cellular quality control pathways. Instead, our results indicate that transient ribosome collisions promote productive translation by suppressing ribosome pausing when ribosomes encounter problems, a process we term ribosome cooperativity. Ribosome cooperativity also suppresses ribosome recycling on problematic sequences, likely by reducing pause duration, thus stimulating processive translation. Together, our results show that ribosomes cooperate during translation to ensure fast and efficient translation.

## Introduction

Fast and accurate translation is essential for all of life, since translation determines the protein content of cells. During mRNA translation, ribosomes catalyze the synthesis of a polypeptide chain based on the sequence of an mRNA. To ensure accurate, error-free protein synthesis, translation is constantly monitored by cellular surveillance mechanisms. Defects in translation regulation, fidelity or quality control can lead to several diseases, including neurodegenerative diseases (D’Orazio and Green, 2021; Knight et al., 2020).

mRNA translation is initiated upon recruitment of the small ribosomal subunit to the 5’ end of the mRNA, followed by 5’UTR scanning, translation start codon recognition and recruitment of the large ribosomal subunit to form a translation-competent 80S ribosome. Following 80S assembly, decoding of the mRNA sequence and synthesis of the polypeptide chain occur during the translation elongation phase. Translation of a single codon consists of multiple steps, including decoding (delivery of a cognate amino-acyl tRNA to the ribosomal A-site), transfer of the growing peptide chain from the P-site peptidyl-tRNA to the A-site aminoacyl-tRNA, a reaction that is catalyzed by the peptidyl transferase center of the ribosome, and finally translocation, during which tRNAs and mRNA are moved relative to the ribosome by exactly one codon (Dever et al., 2018).

During the translation elongation phase, ribosomes can encounter various obstacles that slow down or stall translation. For instance, mRNAs can encode programmed translational pause sites that facilitate correct co-translational folding of the nascent polypeptide, or promote efficient co-translational targeting to an organelle (e.g., the endoplasmic reticulum) (Collart and Weiss, 2020; Zhao et al., 2021). Programmed translational pausing can also allow complex regulation of the mRNA, as is the case for the pause sequence in the Xbp1 mRNA, which is required for efficient mRNA processing during cellular stress (Yanagitani et al., 2011). RNA structures in mRNAs can also slow down translation, which is leveraged by viral RNAs to pause ribosomes at specific sites in their mRNA to induce programmed ribosome frameshifting (Hill and Brierley, 2023; Namy et al., 2006). In addition to *regulated* pauses, ribosome pausing can also occur on defective or damaged mRNA. For example, incorrect pre-mRNA processing can produce mRNAs that are prematurely poly-adenylated, resulting in poly(A) sequences within the coding sequence of an mRNA that potently pause ribosomes (Chandrasekaran et al., 2019). Finally, environmental stresses, such as chemical or UV damage, can chemically modify RNA nucleotides in ways that block translation by ribosomes, resulting in ribosome pausing at sites of mRNA damage (Snieckute et al., 2023; Stoneley et al., 2022; Yan et al., 2019).

Different types of mRNA obstacles slow down ribosomes through different mechanisms. First, pausing can occur through slow decoding of the mRNA. For example, if the cognate tRNA for the A-site codon is expressed at relatively low levels, or if the cognate tRNA pool is mostly uncharged due to an amino acid deficiency in the cell, decoding will be slowed down (Chen and Tanaka, 2018; Gobet et al., 2020). Second, interactions of the nascent polypeptide with the ribosome exit tunnel can also cause ribosome pausing by preventing peptide transfer from the peptidyl-tRNA to the aminoacyl-tRNA, as is the case for some positively charged homopolymeric peptides (e.g., polylysine) (Chandrasekaran *et al*., 2019), as well as for specific peptide sequences (e.g., the Xbp1 arrest peptide) (Yanagitani *et al*., 2011). Peptidyl transferase activity can also be inhibited during translation of polyproline stretches due to the unusual structural configuration of such amino acid sequences (Gutierrez et al., 2013). Finally, the translocation step of the elongation cycle can also be inhibited, for example by the presence of strong RNA structures immediately downstream of the ribosome (Qu et al., 2011).

Much of our knowledge on the translation elongation cycle comes from biochemical and structural studies (Behrmann et al., 2015; Dever *et al*., 2018). As ribosomes are generally assessed individually in such studies, ribosomes are typically thought of as independently operating units. However, mRNA molecules are usually translated as polysomes by multiple ribosomes simultaneously, but it is mostly unknown how different ribosomes translating the same mRNA functionally interact. One example of ribosome-ribosome functional interaction has recently emerged; if a ribosome is stalled, the trailing ribosome can collide with the stalled ribosome, and this collided ‘disome’ is sensed by cellular quality control pathways to remove the stalled ribosome from the mRNA (Juszkiewicz et al., 2018; Meydan and Guydosh, 2021; Simms et al., 2017; Yip and Shao, 2021). In addition to ribosome removal, this quality control pathway results in degradation of the incompletely synthesized nascent chain (Inada, 2017), and, if the number of collided ribosomes in a cell is large, quality control pathways can even activate a cellular stress response (Wu et al., 2020).

While it is clear that ribosome collisions can signal a defect in translation, it is unknown whether all ribosome collisions represent problematic situations involving a stalled ribosome, or whether (transient) collisions between actively translocating ribosomes also occur. If so, it is unclear if and how surveillance mechanisms can distinguish ‘pathological’ collisions, collisions involving a stalled ribosome, from ‘physiological’ collisions between two translocating ribosomes, and only target the former for ribosome recycling. While structural and sequencing-based approaches have uncovered many key aspects governing ribosome collisions, these approaches are not ideally suited to map the kinetic landscape of ribosome-ribosome interactions in cells. Previously our lab and others have developed real-time single-molecule assays to study translation dynamics in living cells (Morisaki et al., 2016; Pichon et al., 2016; Wang et al., 2016; Wu et al., 2016b; Yan et al., 2016), and this technology has already led to important insights into translational quality control as well (Goldman et al., 2021; Hoek et al., 2019). However, these earlier technologies lack robust single-ribosome resolution, hampering precise kinetic measurements of ribosome-ribosome interactions.

Here, we employ our newly developed live-cell translation-imaging method based on Stopless-ORF Circular RNAs (socRNAs; Madern et al., 2024) together with computational modeling to explore the functional relationships between multiple ribosomes translating the same RNA molecule. By loading either single or multiple ribosomes onto socRNAs and visualizing the translation by each ribosome in real time with very high precision, we find that transient ribosome collisions frequently occur during polysomal translation, but that transient ribosome collisions do not result in ribosome recycling. By comparing collisions between two translocating ribosomes with collisions involving a paused ribosome, we find that a combination of ribosome queue length and collision duration (sub-second vs minutes) allow efficient discrimination between physiological and pathological collisions. Surprisingly, we find that transient ribosome collisions can even be productive by suppressing prolonged ribosome pausing during translation, a process we term ‘ribosome cooperativity’. We find that ribosome cooperativity not only enhances translation elongation rates, but also suppresses ribosome recycling by surveillance mechanisms, resulting in fast and highly processive translation.

## Results

To determine if and how ribosomes functionally interact during translation elongation, a method is needed that allows: 1) highly sensitive measurements of translation elongation kinetics, 2) the ability to monitor translating ribosomes for long periods of time, and 3) precise control over the number of ribosomes translating an mRNA molecule. Our recently developed translation imaging method employing Stopless-ORF circular RNAs (socRNAs, Madern et al., 2024) fulfils these requirements. In the socRNA assay one or multiple ribosomes are loaded onto socRNAs and their translation elongation rate is monitored by fluorescent labeling of the growing nascent chain using the SunTag technology, analogous to the single-mRNA translation imaging approach that we and others have developed for visualizing translation of linear mRNAs (Morisaki *et al*., 2016; Pichon *et al*., 2016; Tanenbaum et al., 2014; Wang *et al*., 2016; Wu et al., 2016a; Yan *et al*., 2016). Long-term tracking of individual socRNAs is facilitated by socRNA tethering to the plasma membrane through the nascent chain using the ALFA-Tag peptide-nanobody system (Figure 1A and Video S1). One or more ribosomes are loaded onto the linear precursor RNA of socRNAs. Importantly, no ribosomes are loaded onto the socRNA after circularization, so each socRNA is translated by a constant and well-defined number of ribosomes throughout the experiment (Figure S1A) (Madern et al., 2024). To experimentally determine the number of ribosomes translating each socRNA molecule, nascent polypeptides are released from the socRNAs at the end of each experiment through addition of the translation inhibitor puromycin. The number of nascent chains, which reflects the number of translating ribosomes associated with each socRNA is then scored (Figures 1B-1D and Video S2). Note that in a small number of cases, GFP foci split before the end of the movie due to (premature) release of one or more ribosomes. Such prematurely released ribosomes were included in the analysis of the number of ribosomes per socRNA shown in Figure 1D. To vary the number of ribosomes associated with each socRNA for different experimental settings, we designed socRNAs of different sizes and also introduced an AUG start codon within the 5’UTR of the linear precursor RNA to guide all scanning ribosomes into the socRNA reading frame. Through these socRNA modifications, we obtained a high and a low ribosomal load socRNA (Figure 1D).

**Figure 1.**
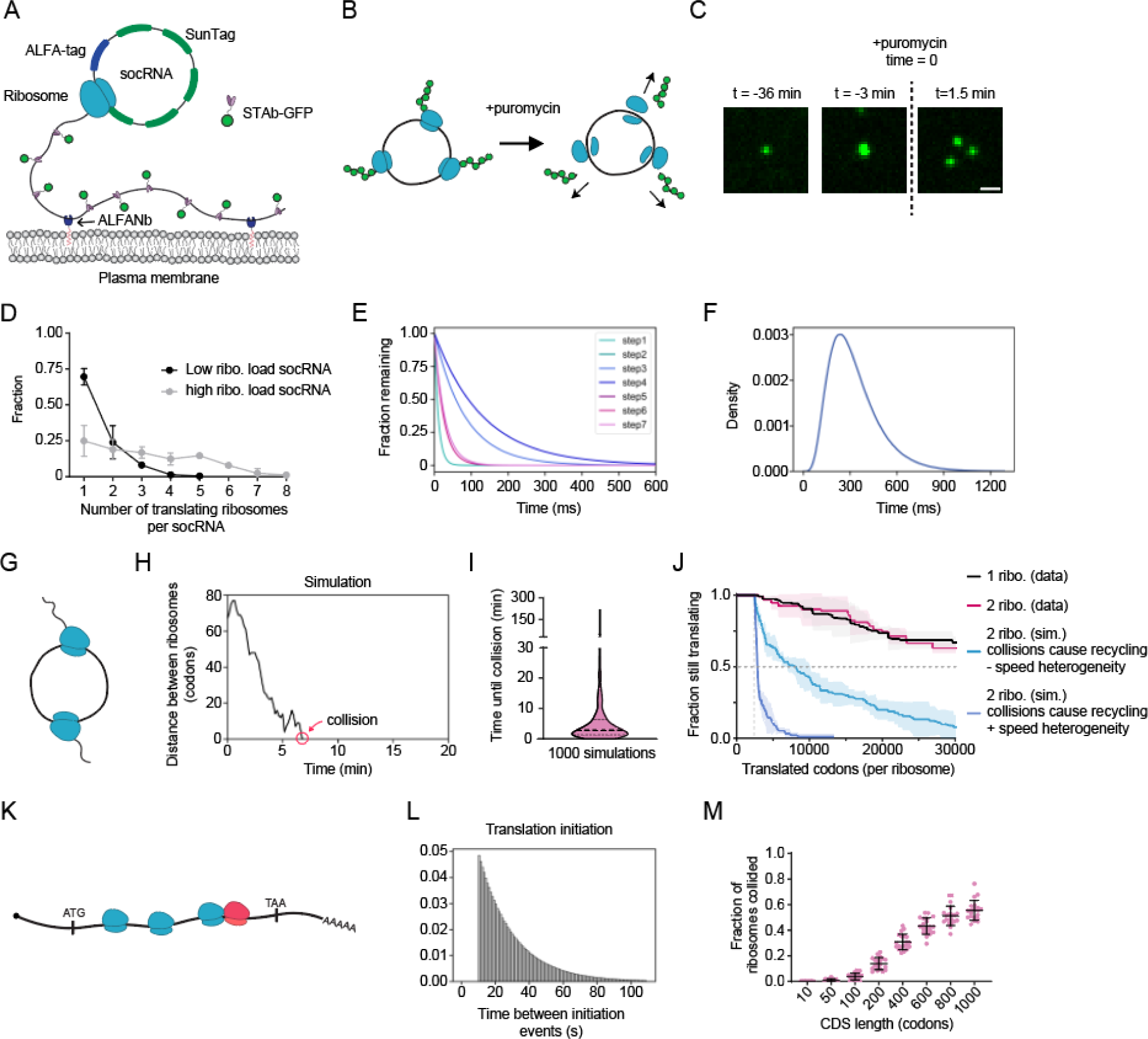
Collisions between translating ribosomes do not induce ribosome recycling. **A**) Schematic of socRNA system. Translation of socRNAs is visualized through binding of GFP-tagged SunTag antibody (STAb-GFP) to nascent SunTag peptides. socRNAs are tethered to the plasma membranes through binding of ALFA-tag peptides to the AlFA-tag nanobody (ALFANb), which is tethered to the plasma membrane. **B**) Schematic showing that socRNAs translated by multiple ribosomes split into multiple GFP foci upon puromycin treatment. **C**) Representative time-lapse images of socRNAs translated by multiple ribosomes (here, 3 ribosomes). Puromycin is added at the indicated time-point, resulting in nascent chain release and splitting of the single GFP spot into three, representing the three polypeptides produced by the three ribosomes translating the socRNA. Scale bar, 1μm. **D**) Distribution of the number of translating ribosomes per socRNA. Two different socRNAs differing in their average ribosomal load were analyzed (black = low ribosome loading, gray = high ribosome loading). Error bars represent standard deviation. **E**) Distributions of the duration of each step in the translation elongation cycle used in simulations. **F**) Distribution of the durations of the complete translation elongation cycle used in simulations. **G**) Schematic depicting socRNA simultaneously translated by two ribosomes. **H**) The time until the first collision between two ribosomes translating the same socRNA was simulated. Initial ribosome positioning on socRNA was randomized. The intrinsic ribosome heterogeneity in elongation speed was incorporated in the simulation. The distance between ribosome A-sites is ploted. Ribosome collisions are called as soon as their A-sites are 10 codons (30 nt) apart. Red circle indicates the moment of collision. **I**) Simulation shown in (H) was repeated 1000 times to determine the median time until 2 ribosomes translating the same socRNA collide. Thick dashed line indicates the median, thin lines indicate 25th and 75th percentile. **J**) U2OS cells stably expressing STAb-GFP, ALFANb-CAAX, and TetR were transfected with socRNAs and imaged by time-lapse microscopy. Kaplan-Meyer survival curve showing the total number of codons translated before aborting translation (referred to as processivity). Black and magenta lines represent experimental data for one or two ribosomes translating a socRNA, respectively. Blue lines indicate data from simulations that include (dark blue) or do not include (light blue) ribosome speed heterogeneity in which transient ribosome collisions result in immediate ribosome recycling. Lines indicate mean values and shaded regions indicate standard deviation. (legend continued on next page) **K**) Cartoon depicting translation of a linear mRNA by multiple ribosomes. The leading ribosome involved in a ribosome collision is highlighted in red. **L**) Distribution of interval between translation initiation events used in simulation shown in (M). **M**) Fraction of ribosomes that undergo at least one collision with a trailing ribosome during translation of the mRNA ploted as a function of mRNA coding sequence (CDS) length. Intrinsic ribosome heterogeneity in elongation speed was incorporated in the simulation. Each data point represents a single simulated mRNA molecule (See methods). Error bars represent standard deviation. The number of experimental repeats and cells analyzed per experiment are listed in Table S1.

Having established an assay that meets the three requirements outlined above for studying functional ribosomal interactions, we first asked whether transient collisions occur between multiple ribosomes translating the same socRNA. Since we recently obtained very precise measurements of the translocation dynamics of individual ribosomes (Madern et al., 2024), we reasoned that we could simulate translation elongation dynamics to determine whether transient collisions occur if socRNAs are translated by more than one ribosome. To build an *in silico* model of translation elongation dynamics, we sub-divided the elongation cycle in seven sub-steps based on structural analysis of the elongation cycle, and determined the average duration of each sub-step based on the relative frequency of occurrences of each sub-step in EM images of unperturbed translating ribosomes (Behrmann *et al*., 2015). Based on these structures, the decoding step and the translocation step are the slowest steps in the elongation cycle and are thus mostly rate-limiting, which qualitatively matches kinetic measurements *in vitro* (Ferguson et al., 2015; Holm et al., 2023; Rodnina et al., 2017). While these sub-step durations represent estimates of elongation cycle kinetics, it is important to note that varying the values for each sub-step of the elongation cycle used in our simulation does not qualitatively affect our conclusions (See below). Stochasticity inherent to all biological processes at the molecular level was introduced by assuming that the duration of each sub-step in the elongation cycle followed an exponential decay distribution, with a mean duration determined as described above (Figure 1E). To simulate ribosome translocation along the mRNA, we randomly drew values from these sub-step time distributions (Figure 1F). We also included the previously observed ribosome elongation speed heterogeneity (Madern et al., 2024) in the simulations (see Methods), and simulations were run 1000 times per condition. As a control, we first simulated elongation dynamics of single ribosomes, and measured the elongation rate per ribosome, which resulted in a very similar ribosome elongation rate distribution as was observed experimentally (compare red bars to black line), confirming that our simulations accurately recapitulate the experimental measurements (Figure S1B). Next, we simulated translation by two ribosomes per socRNA for which ribosome starting positions on socRNAs were randomly selected. Interestingly, two ribosome simulations revealed that translocating ribosomes frequently and rapidly collide on socRNAs, with a median time to collision of just ∼3 minutes (Figures 1G-1I). Since ribosomes typically translate socRNAs for >2 hr, these results show that collisions likely occur frequently on socRNAs. To determine the effect of the selected sub-step duration distributions, we repeated these simulations assuming that one sub-step was very slow (rate-limiting), while the other six were very fast, or assuming that all sub-steps have identical duration distributions. In all cases the average elongation rate was similar. In all cases very similar collision frequencies were observed (Figure S1C-S1F), confirming the validity of these simulations.

We next asked whether collisions between translocating ribosomes cause ribosome recycling. If ribosome collisions between two translating ribosomes on a socRNA induce recycling, the total number of codons translated on a socRNA (referred to here as ‘ribosome processivity’) would be much lower for two ribosomes translating a socRNA compared to one, given the high frequency of collisions identified in our simulations (Figure 1J, light and dark blue lines). We therefore measured ribosome processivity either when ribosomes translated socRNAs on their own, or together with a second ribosome. We transfected a socRNA encoding DNA plasmid into U2OS cells expressing STAb-GFP and a membrane anchored ALFA-tag and performed spinning disk confocal time-lapse imaging of translated socRNAs. We found that ribosome processivity is not affected by the number of ribosomes translating a socRNA (Figure 1J), indicating that transient ribosome collisions do not result in ribosome recycling.

To determine if transient ribosome collisions also occur during translation of linear mRNAs, we simulated translation of linear mRNAs in which we added a translation initiation step to the computational model. We assumed that the time interval between translation initiation events followed an exponential decay distribution with an average interval of 30 sec (a conservative number based on previous measurements (Livingston et al., 2023; Yan *et al*., 2016) and a minimal interval of 10 sec. We then simulated translation on linear mRNAs and determined the frequency of ribosome collisions on mRNAs with different coding sequence lengths (Figures 1K – 1M and Video S3). We find that ribosome collisions also occur frequently on linear mRNAs, with 12% of ribosomes undergoing at least one transient collision during translation of a coding sequence (CDS) of 200 codons and up to ∼60% of ribosomes undergoing one or more collisions on a CDS of 1000 codons. Similar collisions frequencies were observed when using different sub-step duration distributions (Figure S1G). While ribosome elongation speed heterogeneity contributed substantially to the frequent transient ribosome collisions, transient collisions were also observed in the absence of ribosome elongation rate heterogeneity due to the stochastic nature of ribosome translocation (Figure S1H and Video S4). Together, our data support a model in which transient ribosome collisions frequently occur on socRNAs as well as endogenous mRNAs.

Since our results show that transient collisions occur frequently on socRNAs and are not targeted for ribosome recycling, we wondered how such transient, ‘physiological’ collisions are discriminated from persistent, ‘pathological’ collisions, to ensure only the later type is targeted by surveillance mechanisms. One possibility is that physiological collisions are discriminated from pathological collisions based on collision duration. Previous work examined recycling kinetics upon ribosome collision on linear mRNAs using the SunTag-based translation imaging system and a strong pause sequence, which revealed that collided ribosomes are recycled in seconds (Goldman *et al*., 2021). One limitation of previous systems, though, is that mRNAs are typically translated by dozens of ribosomes simultaneously, generating long, steady-state ribosome queues on reporter mRNAs, thus limiting the resolution and quantitative interpretation of ribosome collision and recycling dynamics. To overcome these technical challenges, we devised a set of socRNAs in which collisions between a paused and a translocating ribosome occur under highly controlled conditions.

To induce ribosome pausing, we introduced the well-studied pause sequence of the Xbp1 stress induced transcription factor (Yanagitani *et al*., 2011) into socRNAs (referred to as Xbp1 socRNAs) (Figure 2A). The Xbp1 pause sequence induces ribosomes pausing at a well-defined site, due to an inability of the ribosome to create a peptide bond due to steric effects of the Xbp1 peptide (Shanmuganathan et al., 2019). To further increase the strength of this pause sequence, we also generated a previously described point mutant of the Xbp1 pause sequence (S255A) (Yanagitani *et al*., 2011). To assess pause duration of the Xbp1 wildtype and S255A experimentally, intensities of GFP foci, representing individual translated socRNAs, were quantified over time to assess translation elongation rates. At the end of each experiment, cells were treated with puromycin to quantify of the number of ribosomes translating each socRNA, as described above (See Figure 1B). We first assessed pause duration on socRNAs translated by single ribosomes to determine how long individual ribosomes pause on Xbp1 sequences. As expected, GFP accumulated substantially slower on socRNAs encoding the Xbp1 sequences compared to control socRNAs, with the Xbp1(S255A) sequence showing the strongest effect (Figures 2B-2D). Note that this assay does not allow detection of individual pauses, but rather measures the average duration of translation of the entire socRNA, including the pause. Since the ‘no pause sequence’ control socRNA is identical to the Xbp1 socRNAs except for the pause sequence, the increase in average translation time per full circle for pause sequence containing socRNAs is likely caused by ribosome pausing at the pause site, allowing calculations of pause durations. Importantly, since ribosomes translate pause sequences dozens of times in the socRNA assay, measurements of average pause duration are extremely accurate. Based on the rate of GFP intensity increase, we calculated the pause duration to be 107 sec on the Xbp1(WT) sequence and 337 sec on the Xbp1(S255A) sequence (Figure 2E) (See Methods), confirming the strong pause activity of Xbp1 pause sequences.

**Figure 2.**
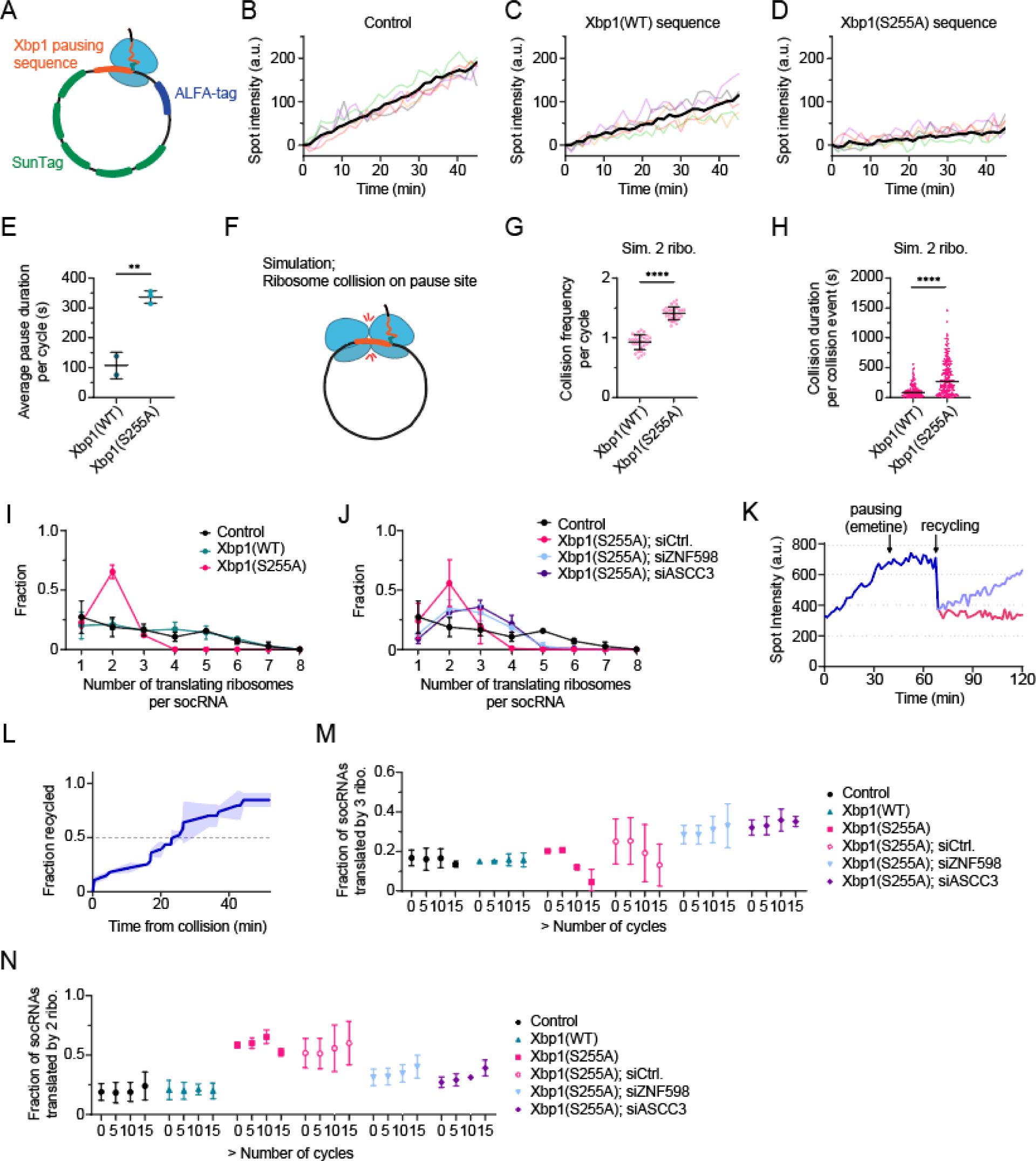
Ribosome collision duration and queue length determine recycling efficiency. **A**) Schematic of socRNA with a ribosome paused on a Xbp1 translation pause sequence. **B-E**) U2OS cells stably expressing STAb-GFP, ALFANb-CAAX, and TetR were transfected with a control socRNA lacking a pause sequence (Control) or with a socRNA encoding indicated Xbp1 pause sequence and imaged by time-lapse microscopy. **B-D**) Representative intensity time traces of single ribosomes translating socRNAs (colored lines). Black line indicates mean values. **E**) Pause durations for each time a ribosome encounters the pause sequence was calculated using GFP intensity time traces for socRNAs translated by a single ribosome (See Methods). Data represent mean ± SD of independent experiments. Dots represent the data from independent experiments. **F-H**) Simulation of ribosome collision at the indicated Xbp1 pause sites on socRNAs translated by two ribosomes. (legend continued on next page) **G**) Collision frequency per cycle at the Xbp1 pause site was calculated from simulations performed for two ribosomes translating socRNAs (See Methods). Note that average pause duration (337 s) is longer than average translation time for one translation cycle (128 s), thus one ribosome can collide twice with the other ribosome during one cycle translation (once as the leading and once as the trailing ribosome), leading to collision frequency per cycle >1. Each dot represents the results from one simulation. Horizontal lines represent mean values ± SD of individual simulation. **H**) Collision durations were calculated from simulations performed for two ribosomes translating Xbp1 socRNAs. Each dot represents an individual collision event. Horizontal lines indicate median values. **I-J**) U2OS cells stably expressing STAb-GFP, ALFANb-CAAX, and TetR were transfected with indicated socRNAs and siRNAs and imaged by time-lapse microscopy. Shown are the distributions of the number of translating ribosomes per socRNAs. Control socRNA (socRNA lacking a pause sequence) data (black dots and line) is reploted from figure 1B for comparison. Data represent mean ± SD of independent experiments. **K-L**) U2OS cells stably expressing STAb-GFP, ALFANb-CAAX, TetR and a control socRNA were followed by time-lapse analysis and GFP intensity of foci was measured over time. A low dose of emetine (3.3 ng/mL) was added to cells at t=30 min to induce collisions between two ribosomes, allowing real-time visualization of collision-induced ribosome recycling. **K**) Representative intensity time trace shows a socRNA translated by two ribosomes, one of which is recycled after a prolonged emetine-induced ribosome collision. **L**) Kaplan-Meyer survival curve showing the time from ribosome collision to recycling for emetine-induced ribosome collisions between two ribosomes. Lines indicate mean values and shaded regions indicate standard deviation. **M-N**) U2OS cells stably expressing STAb-GFP, ALFANb-CAAX, and TetR were transfected with indicated socRNAs and siRNAs and imaged by time-lapse microscopy. Fraction of socRNAs translated by either three (M) or two (N) ribosomes was calculated at different time points. The x-axis represents the estimated number of times the ribosomes has completed translation of a full cycle of the socRNA. Data represent mean ± SD of independent experiments. **, **** indicates p<0.01 and 0.0001, respectively, determined by t-test. The number of experimental repeats and cells analyzed per experiment are listed in Table S1.

Importantly, pause duration on the Xbp1(S255A) socRNA (337 sec) is longer than the time needed for a ribosome to translate a full circle of the socRNA without the pause sequence (128 sec). Therefore, if two or more ribosomes translate Xbp1(S255A) encoding socRNAs, one ribosome will pause long enough for a trailing ribosome to catch up and collide with it in many cases. To obtain quantitative insights into the role of the precise duration of the pause in inducing ribosome collisions, we simulated ribosome translocation and collisions as described in Figure 1, except we added pause sequences to our simulations using experimentally determined pause durations. These simulations revealed that collisions between a paused and a translocating ribosome indeed occur frequently on Xbp1 socRNAs (Figures 2F-2H) and that both collision frequency and collision time were higher for Xbp1(S255A) than Xbp1(WT) socRNAs (Figure 2G and 2H) If collisions between a translating and a paused ribosome on these socRNAs cause ribosome recycling, the number of ribosomes on socRNAs encoding Xbp1 pause sequences should become progressively lower over time compared to control socRNAs, since ribosomes are exclusively loaded on socRNAs at the start of the experiment (Madern et al., 2024). For these experiments, high ribosomal load socRNAs were used (See Figure 1D). When analyzing ribosome number per socRNA after 10 full cycles of translation, the number of ribosomes per Xbp1(S255A) socRNA was strongly reduced compared to control socRNAs (Figure 2I), indicative of collision-induced ribosome recycling. To confirm that the reduction in ribosome load on Xbp1(S255A) socRNA was due to collisions, we depleted ZNF598 and ASCC3 (Figures S2A and S2B), two factors known to be involved in this process (Garzia et al., 2017; Ikeuchi et al., 2019; Juszkiewicz *et al*., 2018; Juszkiewicz et al., 2020; Matsuo et al., 2020). Depletion of either ZNF598 or ASCC3 by siRNA increased the average number of translating ribosomes on Xbp1(S255A) socRNAs (Figure 2J), demonstrating that the reduction of the number of ribosomes on Xbp1 socRNAs indeed reflects collision-induced ribosome recycling. Surprisingly though, while Xbp1(S255A) socRNAs showed a substantial drop in the number of translating ribosomes, the number of ribosomes on Xbp1(WT) socRNAs was indistinguishable from control reporters, even though our simulations show that collisions between a paused and a translocating ribosome occur frequently on Xbp1(WT) socRNAs as well, albeit with shorter duration (Figures 2G and 2H). These results suggest that relatively brief collisions (∼ tens of seconds, figure 2H) between paused and translocating ribosomes are not targeted for ribosome recycling, but when collisions persist for longer periods of time (∼minutes) ZNF598 and ASCC3 sense such collisions and target them for recycling.

Since we cannot identify the exact moment of pausing for Xbp1 socRNAs, measurements of collision duration are only semi-quantitative in those experiments. Therefore, to directly determine the kinetics of ribosome recycling upon collision, we imaged control socRNAs translated by either one or two ribosomes and added the translation inhibitor emetine at a defined moment during the experiment. Using emetine treatment, the precise moment of drug-induced ribosome collision can be assessed based on GFP intensity time traces, as such traces should plateau upon emetine binding to a ribosome (Figure 2K). We titrated emetine concentration to very low levels to ensure that only one ribosome per socRNA was targeted by the elongation inhibitor, allowing the other ribosome to collide with the emetine-arrested ribosome (Figure S2C). To prevent overloading of the cell-wide ribosome collision surveillance pathway upon low-dose emetine addition, harringtonine was added to cells 30 minutes prior to emetine addition, which results in ribosome run-off on endogenous linear mRNAs, but not on socRNAs. Addition of emetine to socRNAs translated by multiple ribosomes resulted in occasional plateaus in GFP intensity time traces, followed by ribosome splitting, and, frequently, resumption of translation by the remaining ribosomes (Figure 2K). To calculate the time from ribosome collision to recycling, we fit a two-state linear model to GFP intensity time traces to determine the precise moment of plateauing (Figure S2D). To call the moment of collision, we computationally correct for the average time needed for a translating ribosome to arrive at and collide with the emetine-arrested ribosome on the same socRNA (∼0.5 min). To precisely call the moment of ribosome recycling, we determined the moment of visible GFP foci splitting, which we corrected for the average duration between ribosome recycling and visible GFP foci separation (i.e., the time needed for ribosome nascent chains to diffuse apart from each other), as assessed by puromycin-induced nascent chain release (∼2 min, Figure S2E). After incorporating these corrections, our results revealed that ribosome recycling upon collision is surprisingly slow, with a half-life of 22 minutes from collision until ribosome recycling (Figure 2L).

A second interesting finding from abovementioned ribosomal load measurements on control and Xbp1 socRNAs is that many (∼70 %) Xbp1(S255A) socRNAs were still translated by *two* ribosomes after 10 cycles of translation, even though the number of socRNAs with *three* or more ribosomes was strongly reduced. If prolonged collisions (∼minutes) between two ribosomes results in effective ribosome recycling, all Xbp1(S255A) socRNAs should be translated by a single ribosome after >10 full cycles of translation, since the two ribosomes will have undergone many prolonged collisions. These results suggest that collisions between many ribosomes (i.e., long ribosome queues) cause efficient ribosome recycling, while collisions between two ribosomes are recycled less efficiently. To examine the impact of ribosome queue length on ribosome recycling kinetics more quantitatively, we examined the number of ribosomes per socRNA at different time points after initial ribosome loading. While on control and Xbp1(WT) socRNAs the number of socRNAs containing three ribosomes remained constant over time, this number was reduced over time for Xbp1(S255A), an effect that was dependent on ZNF598 and ASCC3 (Figure 2M), confirming that this reduction was caused by ribosome recycling. Interestingly, it took ∼10 cycles of translation, and thus ∼10 prolonged collisions, to reduce the number of socRNAs translated by three ribosomes by just 50%, indicating that even at average collision durations of ∼5.6 minutes, recycling efficiency is relatively low (<10%), consistent with our independent result that recycling takes on average ∼20 min upon ribosome collision induced by emetine (Figure 2L), indicating that the calculated recycling time was consistent among different types of ribosome stalls. In contrast to the number of Xbp1(S255A) socRNAs translated by three ribosomes, the number of socRNAs translated by two ribosomes was relatively constant over time (Figure 2N), further suggesting that collisions between two ribosomes may be recycled less efficiently than those between ribosomes in longer queues. In summary, these results suggest that collision duration and queue length together shape recycling efficiency. Since both collision duration and queue length will be longer for collisions with a stalled ribosome compared to transient collisions between two translocating ribosomes, these results explain how these two types of collisions can be efficiently discriminated.

Our results show that transient collisions between ribosomes do not generally result in ribosome recycling, even when one ribosome is briefly paused (e.g., on the Xbp1(WT) sequence), raising the question whether such transient collisions effect ribosome translocation in other ways. To understand the effects of ribosome collisions on ribosome translocation dynamics, we turned to computational simulations again. Interestingly, simulating socRNAs translated by either one or two ribosomes revealed that ribosomes move slightly slower when translating an RNA together with a second ribosome (Figure 3A), an effect we term ‘ribosome interference’. Such ribosome interference is expected as a result of heterogeneous ribosome translocation speeds and the inability of a faster ribosome to overtake the slower one. While ribosome interference is minimal when both ribosomes are continuously translocating, our simulations showed that the effect becomes more pronounced when a pause site is introduced into a socRNA (Figure 3B and Movie S5), since a paused ribosome blocks translocation of the trailing ribosome. To test for ribosome interference experimentally, we measured ribosome pause duration at Xbp1(S255A) pause sites in the case where socRNAs are translated by either one or two ribosomes. We focused on ribosome interference in the presence of ribosome pausing, as the effects of ribosome interference in the absence of pause sites is likely too subtle to detect experimentally. Strikingly, and in contrast to our simulations, the experimentally determined pause duration for two ribosomes translating the same Xbp1(S255A) socRNA was substantially shorter than expected based on simulations of ribosome interference (Figures 3C and 3D). In fact, the average pause time per ribosome on a Xbp1(S255A) socRNA translated by two ribosomes was even shorter than the pause time of single ribosomes translating the same sequence (Figures 3C and 3D). In contrast, on control socRNAs, translation elongation rates were similar for socRNAs translated by one or two ribosomes (Figure 3A), suggesting that the second ribosome specifically suppresses pausing on the pause site, rather than (substantially) speeding up translation on normal mRNA sequences. Together, these results reveal that ribosomes can help each other overcome strong pauses, a mechanism we refer to as ‘ribosome cooperativity’.

**Figure 3.**
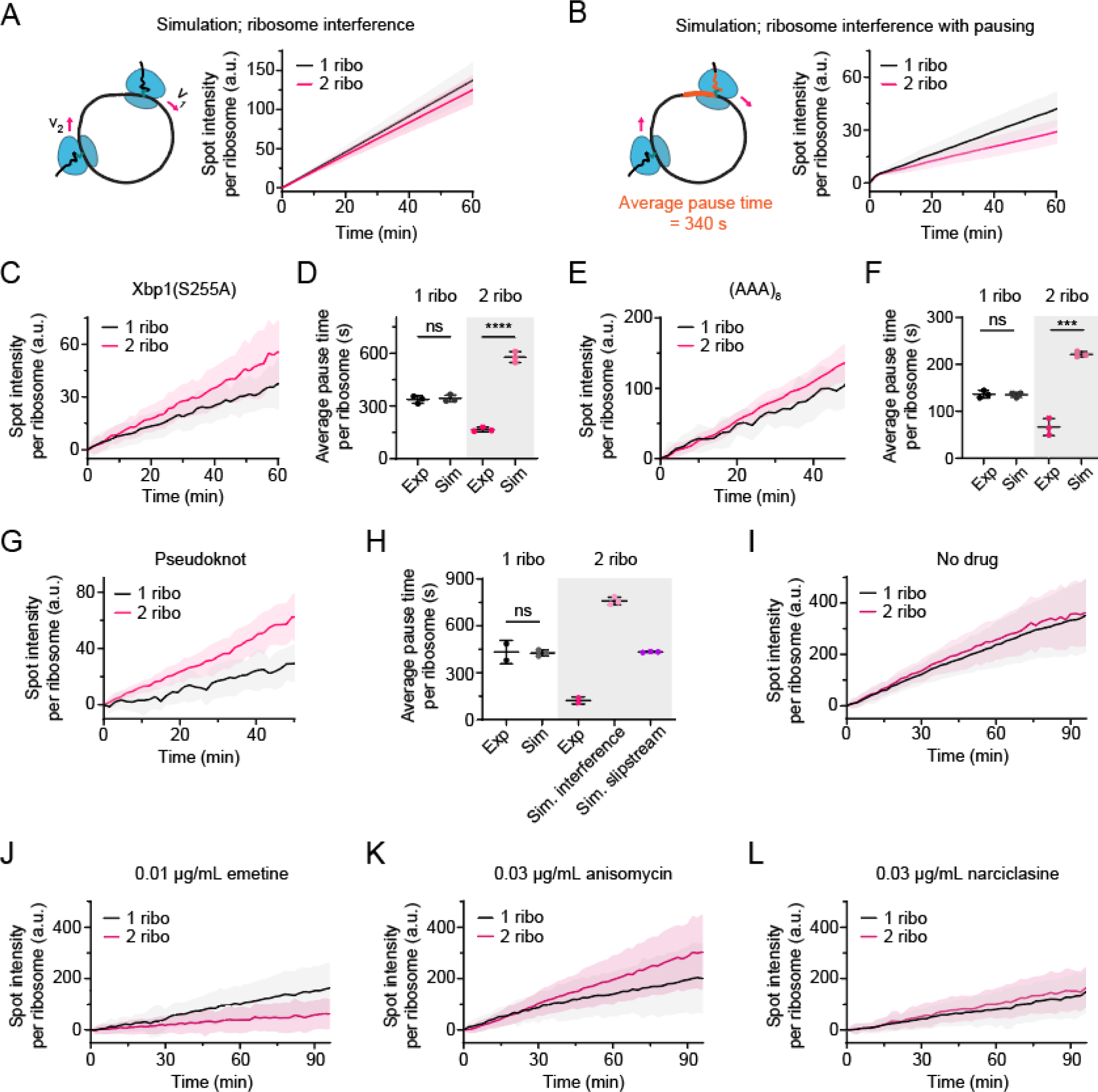
Ribosome cooperativity suppresses ribosome pausing. **A-B**) Simulation of translation on socRNAs with (A) or without (B) pause site. GFP foci intensity per ribosome over time was calculated for single ribosome (black) and two ribosomes (magenta) translating the same socRNA (See Methods). **C-H**) U2OS cells stably expressing STAb-GFP, ALFANb-CAAX, and TetR were transfected with indicated socRNAs and imaged by time-lapse microscopy. Inserted pause sequence: (C-D) Xbp1(S255A), (E-F) (AAA)_8,_ (G-H) RNA pseudoknot. **C, E, G**) Representative intensity time traces of single (black) or two (magenta) ribosomes translating socRNAs. Lines indicate mean values and shaded regions indicate standard deviation. **D, F, H**) Pause durations for each time a ribosome encounters a pause sequence was calculated for both experiments and simulations. Black horizontal lines represent mean ± SD of independent experiments or simulations and each dot represents the data from one independent experiment or simulation. **I-L**) U2OS cells stably expressing STAb-GFP, ALFANb-CAAX, and TetR were transfected with socRNAs and imaged by time-lapse microscopy. Cells were treated with indicated low concentrations of elongation inhibitors to determine the effect of drug-induced pausing on ribosomes translating a socRNA either alone or together with a second ribosome. socRNA spot intensities were measured over time and normalized to the number of ribosomes on the socRNA for direct comparison (see Methods). Lines indicate mean values and shaded regions indicate standard deviation. *, **, ***, **** indicate p<0.05, 0.01, 0.001, and 0.0001, respectively, determined by t-test. The number of experimental repeats and cells analyzed per experiment are listed in Table S1.

We wondered whether ribosome cooperativity could overcome other types of translation pauses as well. First, we assessed the pause time on socRNAs encoding a stretch of 24 A’s, which is known to slow down ribosome translocation (Arthur et al., 2015a; Chandrasekaran *et al*., 2019; Goldman *et al*., 2021). Similar to the Xbp1(S255A) socRNAs, the average pause duration for two ribosomes translating (AAA)_8_ socRNAs was substantially shorter than expected based on simulations (which included ribosome interference) and even somewhat faster than single ribosomes translating the same sequence (Figures 3E and 3F). We also examined socRNAs containing an RNA pseudoknot, a very strong RNA structure (Niu et al., 2021) that acts as a strong pause site for translocating ribosomes (Figures 3G and 3H). Similar to Xbp1(S255A) and poly(A) sequences, two ribosomes paused at RNA pseudoknots far shorter than expected based on simulations and also shorter than single ribosomes (Figures 3G and 3H). Together, these results show that ribosome cooperativity suppresses ribosome pausing on a variety of different pause sequences. For RNA pseudoknot socRNAs, we considered an several explanations for the apparent ribosome cooperativity; if one ribosome pauses for a prolonged period of time upstream of the RNA structure, the second ribosome will catch up with the leading one. Once the first ribosome has successfully unfolded the structure and starts translating the RNA structure sequence, the trailing ribosome will follow closely, preventing the structure from reforming in between the two ribosomes. In this scenario, the trailing ribosome is not impeded by the RNA structure due to the presence of the leading ribosome, somewhat analogous to a ‘slipstream’ effect. To test whether the slipstream effect is sufficient to explain shorter pause times for two vs one ribosome on pseudoknot socRNAs, we included a slipstream effect in our simulations. While the slipstream effect reduced the expected pause time per ribosome for socRNAs translated by two ribosomes compared to one, experiments showed that two ribosomes still moved substantially faster than expected for the slipstream effect alone, indicating that ribosome cooperativity also suppresses pause duration for pseudoknot socRNAs (Figure 3H).

Finally, we asked whether drug-induced ribosome pauses could also be suppressed by ribosome cooperativity. To this end, we treated cells with low doses of either emetine, anisomycin or narciclasine, three translation elongation inhibitors, and measured average translation rates on socRNAs (ploted as the translation elongation rate per ribosome). In the absence of ribosome cooperativity, translation rates per ribosome should be reduced by ∼50% if a socRNA is translated by two ribosomes, since the probability of one of the two ribosomes on the socRNA binding a drug molecule is twice as high compared to socRNAs translated by a single ribosome. Indeed, for low dose emetine treatment, we find that average translation rates per ribosome are reduced by ∼50% when comparing socRNAs translated by one vs two ribosomes (Figures 3I and 3J), indicating that ribosome cooperativity does not act on emetine-bound ribosomes. However, for both anisomycin and narciclasine, translation rates for socRNAs translated by two ribosomes was substantially higher than expected, indicative of ribosome cooperativity (Figure 3K and 3L). Interestingly, both anisomycin and narciclasine likely inhibit the step of peptide bond formation in the translation elongation cycle (Garreau de Loubresse et al., 2014), similar to the Xbp1 sequence (Shanmuganathan *et al*., 2019) and the polylysine peptide encoded by the poly(A) sequence (Chandrasekaran *et al*., 2019) (although poly(A) sequences also inhibit the decoding step (Tesina et al., 2020)), while emetine inhibits the mRNA-tRNA translocation step (Eiler et al., 2024; Wong et al., 2014), suggesting that ribosome cooperativity may promote peptide bond formation. Additional steps in the elongation cycle may also be facilitated by ribosome cooperativity, as RNA structures, which are also targeted by ribosome cooperativity, likely inhibit the translocation step of the elongation cycle (Caliskan et al., 2014; Namy *et al*., 2006)(See Discussion). Together, these results show that ribosome cooperativity helps resolve prolonged translational pauses, likely through multiple mechanisms.

Next, we asked how ribosomes could cooperate to overcome translational pauses. We considered a model in which a translocating ribosome ‘pushes’ the paused ribosome upon collision to resolve the pause and to stimulate resumption of translocation of the leading ribosome. We simulated socRNA translation dynamics assuming that collisions instantaneously resolved the pause upon initial collision (Figures 4A and 4B, Video S6). For both the Xbp1(S255A) and the poly(A) reporter, these simulations showed that collision-induced pause resolution explains the reduction in pause duration well, although simulations predicted a slightly stronger pause reduction than was observed in experiments. Addition of either a small delay (∼15-40 sec) between collision and resumption of translation to simulations, or including the parameter that not all collision events successfully suppress pausing both allowed quantitative matching of simulations and experiments (Figures S3A-S3D).

**Figure 4.**
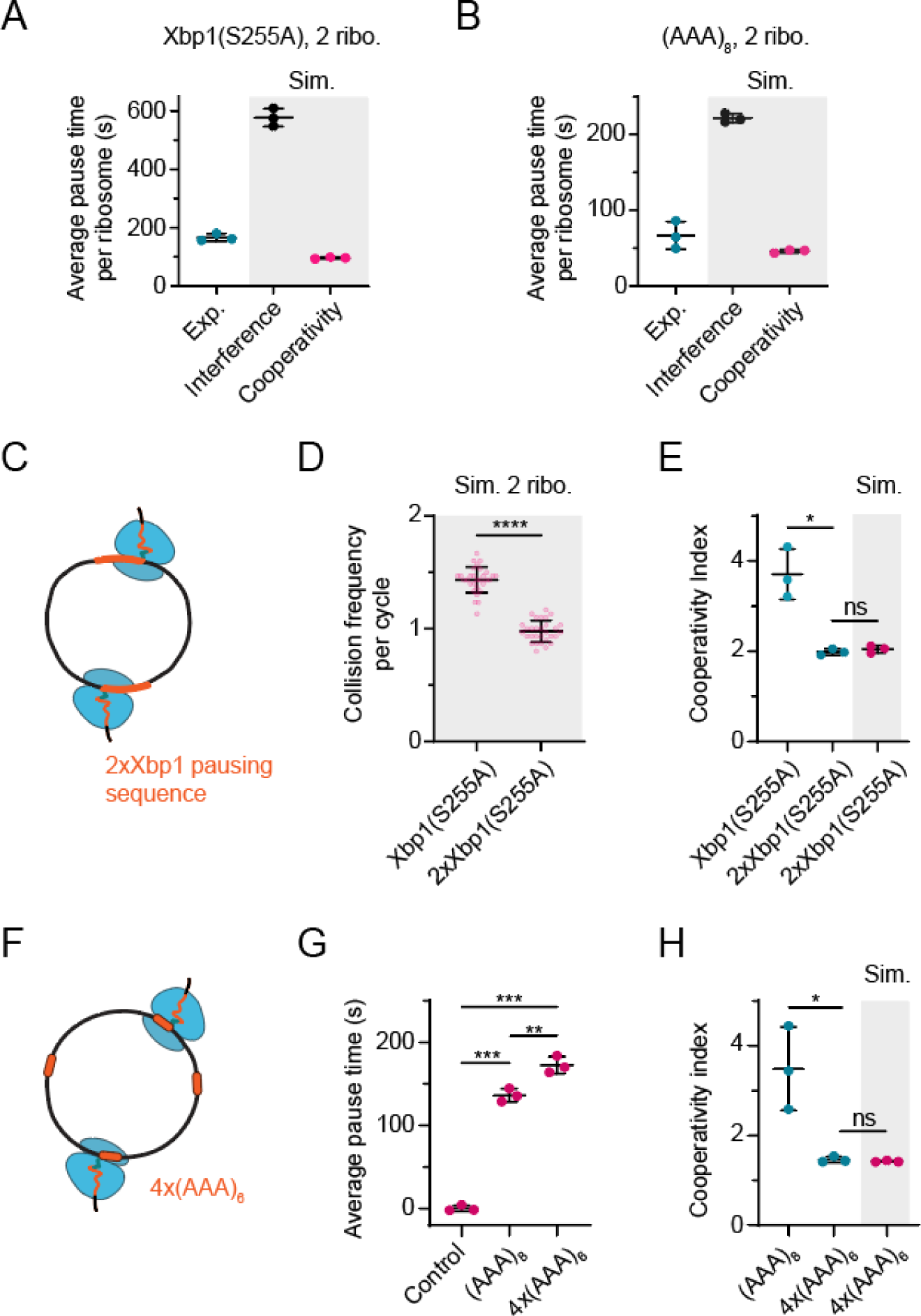
Evidence that collisions underlie ribosome cooperativity. **A-B**) Simulation of average pause time on Xbp1(S255A) (A) or (AAA)_8_ (B) pause sequences for socRNAs translated by two ribosomes. Collision-induced resumption of translation of the paused ribosome were simulated for cooperativity model. Cyan (experimental data) and black (interference model) dots are reploted from Figure 3D and Figure 3F for comparison. **C**) Schematic depicting a socRNA containing two Xbp1 pause sequences introduced at different locations in a socRNA translated simultaneously by two ribosomes. **D**) Collision frequency per full cycle of translation of the socRNA was calculated using simulations. Note that we counted collision events at only one of the two pause sites for 2xXbp1(S255A) socRNA. **E**) The cooperativity index for socRNAs containing either a single or two Xbp1 pause sequences was calculated from simulations (magenta) and experiments (cyan) (See Methods). The cooperativity index represents a measure for the degree to which the presence of a second ribosomes speeds up the average translation elongation speed of a ribosome (See Methods). Simulations quantitatively match the experimentally observed decrease in ribosome cooperativity upon introduction of a second Xbp1 pause sequence. **F**) Schematic depicting socRNA with four (AAA)_6_ pause sequences introduced at different locations within the socRNA. Two ribosomes are simultaneously translating the socRNA. **G**) The average pause duration for indicated socRNAs is shown. Note that the sum of all four pauses is shown for the 4x(AAA)_6_ socRNA. **H**) The cooperativity index for socRNAs containing a single (AAA)_8_ or four (AAA)_6_ was calculated. Simulations quantitatively match the experimentally observed decrease in ribosome cooperativity upon replacing a single, strong (AAA)_8_ pause sequence with four weaker, spatially separated (AAA)_6_ pause sequences. (legend continued on next page) *, **, ***, **** indicate p<0.05, 0.01, 0.001, and 0.0001, respectively, determined by t-test. The horizontal black lines and error bars represent mean ± SD of independent experiments or simulation. The dots represent the data from independent experiments or simulations. The number of experimental repeats and cells analyzed per experiment are listed in Table S1.

To experimentally validate that ribosome collisions underlie ribosome cooperativity, we designed perturbation experiments in which a socRNAs were generated with reduced ribosome collision frequency. To reduce collision frequency, a second Xbp1(S255A) pause sequence was introduced in the Xbp1(S255A) socRNA (Figure 4C). As expected, simulations showed that ribosome collisions occur less frequently on socRNAs containing two pause sequences instead of one (Figure 4D), because both ribosomes are paused at opposite sides of the socRNA for a substantial fraction of the total time, during which time they cannot collide. If ribosome collisions indeed underlie ribosome cooperativity, then socRNAs with multiple pause sites should show less ribosome cooperativity than socRNAs with a single pause site, a hypothesis that was confirmed using simulations (Figure 4E). When we experimentally determined translation rates on socRNAs containing either one or two Xbp1(S255a) pause sequences, we found that ribosome cooperativity was indeed reduced by introduction of the second pause site, and the magnitude of this effect was almost identical to our simulations (Figure 4E). To further validate the importance of collisions for ribosome cooperativity, we also introduced multiple pause sequences for poly(A) reporters to reduce collision frequency. We generated poly(A) reporters containing either a single (AAA)_8_ pause sequence or four shorter (AAA)_6_ which together had a total pause duration that was similar to the single longer poly(A) stretch (136 vs. 173 seconds, Figure 4G). Similar to the 2xXbp1 socRNAs, we found that cooperativity was reduced by introduction of multiple poly(A) sequences to the extent predicted by our simulations (Figure 4H). Together, these results strongly suggest that collisions drive ribosome cooperativity, likely by suppressing ribosome pausing at problematic sequences.

socRNAs not only allow precise measurements of translation elongation rates, but also of ribosome processivity, the total number of codons translated by the ribosome before translation is aborted. Since ribosome cooperativity suppresses pausing, we asked whether processivity was also altered by ribosome cooperativity. We measured the total number of codons translated per ribosome when socRNAs were translated by either one or two ribosomes (See Methods). In control socRNAs, processivity was similar for socRNAs translated by one or two ribosomes (See Figure 1J). Introduction of either a Xbp1(S255A) sequence, a pseudoknot RNA structure or a poly(A) sequence reduced processivity substantially when socRNAs were translated by one ribosome (Figures S4A-S4C). However, ribosome processivity was strongly enhanced when pause reporters were translated by two ribosomes (Figures 5A-5C), almost completely rescuing the decrease in processivity caused by the pause sequences (Figures S4D-S4E). Thus, we conclude that ribosome cooperativity enhances both translation speed and processivity.

**Figure 5.**
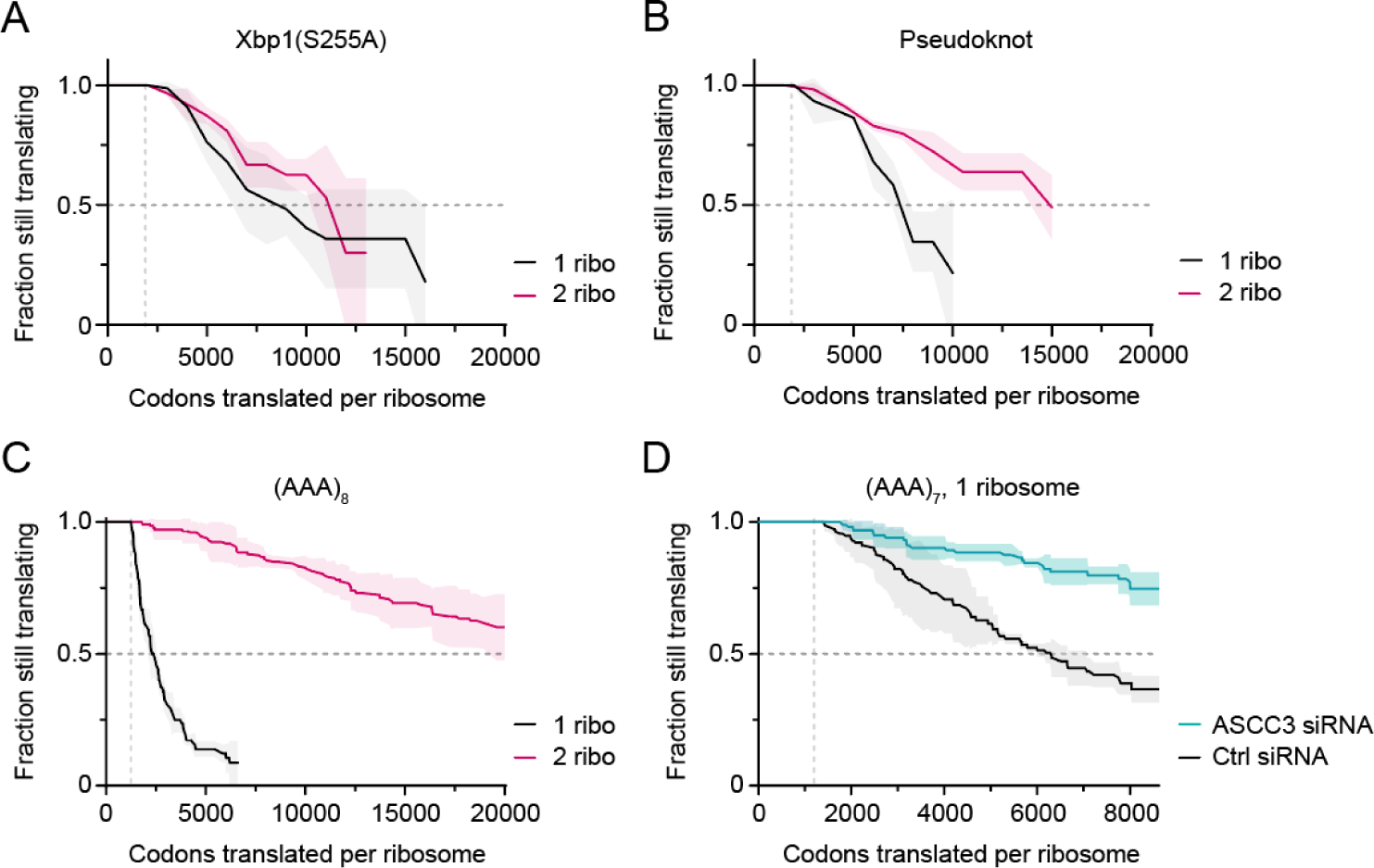
Ribosome cooperativity enhances processivity by suppressing ribosome recycling. **A-D**) U2OS cells stably expressing STAb-GFP, ALFANb-CAAX, and TetR were transfected with indicated socRNAs (A-D) and indicated siRNAs (D) and imaged by time-lapse microscopy. Kaplan-Meyer survival curves of indicated socRNAs show the total number of codons translated by ribosomes before aborting translation. Lines indicate mean values and shaded regions indicate standard deviation. The number of experimental repeats and cells analyzed per experiment are listed in Table S1.

How could ribosome cooperativity stimulate processivity on pause reporters? To answer this question, we need to understand why single ribosomes translating a socRNA containing a pause site have a reduced processivity, as ribosome cooperativity likely suppresses this mechanism. Ribosome processivity can, in principle, be affected by a number of different processes, including premature termination, frameshifting, socRNA cleavage, and ribosome recycling after triggering of ribosome surveillance mechanisms. Earlier work has mainly implicated ribosome collisions in inducing ribosome recycling on pause sites (Juszkiewicz *et al*., 2020; Matsuo *et al*., 2020). A recent study in yeast suggested that single stalled ribosomes may also be targeted by surveillance mechanisms (Li et al., 2022), but no such pathway has been identified in human cells. Our socRNA approach uniquely allows analysis of ribosome recycling on pause sites in the absence of ribosome collisions (i.e., when socRNAs are translated by a single ribosome). To determine whether the decreased processivity of single ribosomes on pause site containing socRNAs was caused by ribosome surveillance mechanisms, we depleted the helicase ASCC3, which is responsible for ribosome recycling downstream of translation surveillance mechanisms (Juszkiewicz *et al*., 2020; Matsuo *et al*., 2020). Indeed, when examining ribosome processivity on the poly(A) socRNA, we found that processivity was largely restored upon depletion of ASCC3, indicating that the reduction in processivity on these socRNAs was indeed due to triggering of quality control pathways by single paused ribosomes (Figure 5D). Thus, we identify a pathway in human cells that efficiently targets single, paused ribosomes for recycling, and our results suggest that ribosome cooperativity enhances processivity by suppressing the surveillance pathway targeting single ribosomes.

We recently found that ribosomes undergo stochastic pauses on ‘normal’ mRNA sequences (i.e., those that don’t encode pause sequences) (Madern et al., 2024). We therefore asked whether ribosome cooperativity also helps to overcome such stochastic pauses. To identify stochastic pauses during socRNA translation, we measured GFP intensities over time and identified stochastic pauses in intensity time traces using Hidden Markov Modeling (Figures 6A and 6B and see Methods). We analyzed GFP intensity time traces of socRNAs translated by either one or ≥two ribosomes. We could identify prolonged pauses in ∼5% of intensity time traces of socRNAs translated by single ribosomes. We next examined ribosome pausing for socRNAs translated by two or more ribosomes. Importantly, assuming that each ribosome has an equal probability of pausing during translation, more pauses are expected in GFP intensity time traces for socRNAs translated by multiple ribosomes, as a (prolonged) pause of one of the ribosomes blocks elongation of other ribosomes on the socRNA resulting in a plateau in the intensity time trace. However, we did not detect any pauses for socRNAs translated by two or more ribosomes, suggesting that ribosome cooperativity indeed suppresses stochastic ribosome pausing as well (Figure 6C). To exclude that the absence of detectable pauses in the case of socRNAs translated by multiple ribosomes was due to rapid recycling of paused ribosomes upon collisions, we knocked down ASCC3 (which prevents ribosome recycling) and repeated the pause detection experiments (Figure 6D). Even after ASCC3 knockdown no pauses could be detected when socRNAs were translated by multiple ribosomes. These results show that ribosome cooperativity also acts to suppress stochastic pausing of ribosomes on normal mRNA sequences.

**Figure 6.**
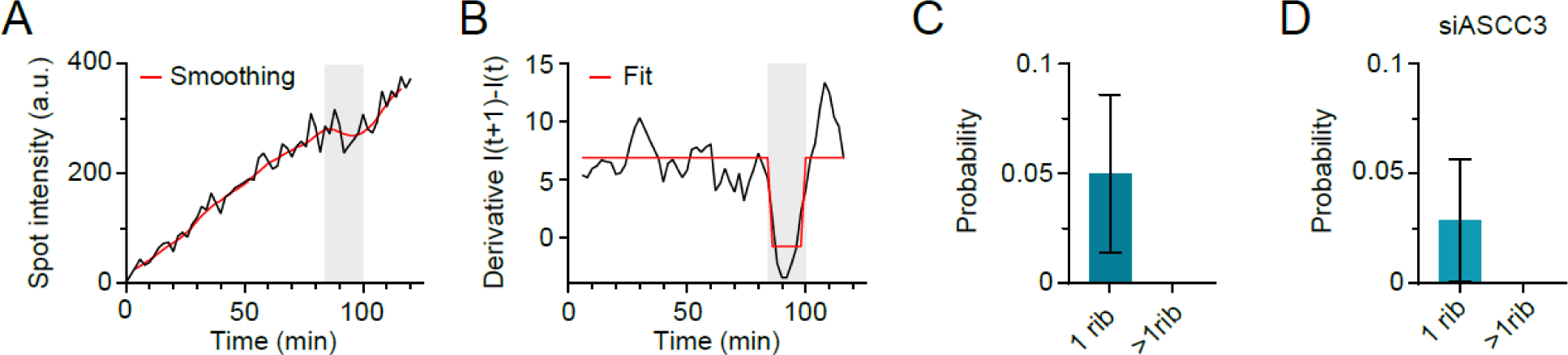
Ribosome cooperativity suppresses stochastic pauses on non-problematic mRNA sequences. **A-D**) U2OS cells stably expressing STAb-GFP, ALFANb-CAAX, and TetR were transfected with socRNAs encoding 5xSunTag and ALFA-tag without pause sequence (A-C) and indicated siRNA (D), and imaged by time-lapse microscopy. **A-B**) Example intensity time trace used for pause identification. **A**) Representative raw intensity time trace (black line) and smoothed data (red line, see Methods). **B**) Derivative of the smoothed example line in (A) (Black line). Red line shows Hidden Markov Modeling to identify translation pauses. **C-D**) Probability of identifying a pause in intensity time traces of socRNAs using the approach shown in (A, B). Error bars indicate standard deviation from independent experiments. The number of experimental repeats and cells analyzed per experiment are listed in Table S1.

## Discussion

In this work, we use a combination of experiments and computer simulations to study how ribosomes interact with each other during translation elongation, an important question that has remained largely unexplored. First, we find that two types of ribosome collisions can occur during translation; The first type, which we call ‘physiological collisions’, are transient collisions between two translocating ribosomes that do not need to be removed from the mRNA. The second type of collisions, ‘pathological collisions’, are collisions between a translocating ribosome and a stalled ribosome, which do require removal of the stalled ribosome for productive translation to resume on the mRNA. Our results suggest that these two types of collisions are likely discriminated based on collision duration and queue length. Surprisingly, we find that transient ribosome collisions can suppress ribosome pausing and reduce recycling to stimulate fast and processive translation. We term this behavior ‘ribosome cooperativity’, and show that ribosome cooperativity occurs both on problematic mRNA sequences and during stochastic ribosome pauses on normal RNA sequences. Together, this work sets a new paradigm for ribosome-ribosome interactions during mRNA translation, and suggests that the high ribosome density on mRNAs typically observed during eukaryotic translation might provide advantages for translation efficiency beyond an increased protein output.

### Discriminating physiological from pathological collisions

The pathways involved in sensing and clearing collided ribosomes have been intensely studied in the past few years, typically by using (strong) pause sequences (Juszkiewicz *et al*., 2018; Juszkiewicz *et al*., 2020; Meydan and Guydosh, 2021; Simms *et al*., 2017; Sundaramoorthy et al., 2017). However, the extent to which ribosome collisions occur on normal mRNA sequences, and whether ribosomes in such collisions are also recycled remained poorly understood. By combining experiments and simulations we show that ribosome collisions also occur between actively translating ribosomes in the form of transient collisions, which are caused both by heterogeneity in translation speeds by different ribosomes, but also by the inherent stochasticity of the translation elongation cycle. Intriguingly, we find that transient collisions, in contrast to pathological collisions, do not induce ribosome recycling. These findings raise the question how such transient collisions are discriminated from pathological collisions by surveillance mechanisms to ensure only ribosomes involved in pathological collisions are sensed and cleared. To address this question, a recent study employed a single-molecule imaging approach using linear mRNAs to measure ribosome recycling kinetics after ribosome collisions (Goldman *et al*., 2021). Somewhat surprisingly, this study found that collision-induced ribosome recycling takes only ∼8 s. While this value is slower than the expected duration of transient collisions, and thus could suggest a mechanism by which transient and persistent collisions are discriminated, the observed a recycling time of ∼8 s contrasts with our recycling rate measurements of ∼20 min. This discrepancy may be caused by a key difference in the experimental setup; the earlier study measured recycling rates of ribosomes in long, steady-state queues of collided ribosomes, while the socRNA approach allowed us to measure the precise time from collision to recycling for individual pairs of collided ribosomes. It is known that collisions induce ribosome subunit ubiquitination, followed by ribosome splitting (Juszkiewicz and Hegde, 2017; Juszkiewicz *et al*., 2020; Matsuo et al., 2017; Sundaramoorthy *et al*., 2017). Possibly, ribosome ubiquitination upon collision is relatively slow (minutes), while splitting of ubiquitinated ribosomes is fast (seconds). In the long steady-state queues in the earlier work, ribosomes may have had sufficient time to become fully ubiquitinated, so the observed recycling kinetics are driven mainly by the fast ribosome splitting rates. In contrast, in our socRNA assay, the time from initial collision to recycling is assessed, thus including both the ubiquitination and ribosome splitting kinetics, resulting in far slower measured recycling times. Importantly, the very slow ribosome recycling rate measured here implies that transient collisions can be robustly discriminated from persistent collisions based on collision duration.

SocRNAs also allow control of the number of ribosomes involved in a collision, which provided evidence that recycling efficiency is also dependent on ribosome queue length (Figure 2). Consistently, a recent *in vitro* study has shown that ubiquitination of collided ribosomes occurs more efficiently in a queue of three vs two ribosomes (Juszkiewicz *et al*., 2018; Matsuo et al., 2023), and even in bacteria there is evidence that longer queues might be targeted by surveillance mechanisms more efficiently (Saito et al., 2022). Mechanistically, longer ribosome queues may cause increased local concentration of surveillance proteins around stalled ribosomes, resulting in a higher rate of ribosome ubiquitination and/or recycling. Together, these observations lead to a unifying model in which short-lived ribosome collisions, which likely occur frequently on mRNAs, are insufficient to induce ribosome recycling and can even relieve ribosome pausing, but if ribosome pausing persists after collision, long and stable ribosome queues will form, resulting in efficient recycling of the stalled ribosome.

### Molecular mechanisms underlying ribosome cooperativity

Our data suggest at least three, non-mutually exclusive mechanisms underlying ribosome cooperativity, which may act to resolve different types of ribosome pauses. First, for pauses on highly structured RNAs, we propose that a ‘slipstream’ effect speeds up translation in the presence of multiple ribosomes. If two ribosomes are very close together during translation and encounter a strong RNA structure, the structure only needs to be unfolded once (by the leading ribosome), instead of twice (by both ribosomes), to allow both ribosomes to pass the structure, as the structure cannot reform in between the two closely adjacent translocating ribosomes. This magnitude of this slipstream effect is, however, not sufficient to explain the cooperativity of ribosomes translating a structured RNA in our assays (Figure 3H), suggesting that a second type of cooperativity must exist. We speculate that two adjacent ribosomes can simultaneously pull on the mRNA to produce more force than a single ribosome to unwind the upstream RNA structure. Third, we find that ribosome cooperativity also suppresses pausing when peptide bond formation is inhibited (e.g., Xbp1 pause sequence and pauses induced anisomycin and narciclasine, Figure 3) (Dmitriev et al., 2020; Garreau de Loubresse *et al*., 2014; Shanmuganathan *et al*., 2019), suggesting that peptide bond formation may be directly or indirectly stimulated by ribosome collisions. Peptide bond formation is also inhibited by poly(A) sequences through the formation of highly basic polylysine peptides (Arthur et al., 2015b; Chandrasekaran *et al*., 2019). However, poly(A) sequences also induce mRNA structural defects in the decoding center (Chandrasekaran *et al*., 2019; Tesina *et al*., 2020), so it is possible that either one, or both steps are stimulated by ribosome cooperativity on poly(A) sequences. Intriguingly, ribosome cooperativity also suppresses stochastic pauses during translation of non-problematic sequences, and it will be interesting to study whether such stochastic pauses are caused by RNA structures or peptide bond formation defects as well. Future studies will hopefully also shed further light on the structural mechanisms underlying ribosome cooperativity during translation of problematic and non-problematic mRNA sequences.

### Ribosome cooperativity suppresses translation surveillance mechanisms

Previous work has established that ribosome collisions are used as a proxy for sensing ribosome stalling during mRNA translation, leading to the recruitment of the cellular quality control machinery to recycle the stalled ribosome (Ikeuchi *et al*., 2019; Juszkiewicz *et al*., 2018; Juszkiewicz *et al*., 2020; Meydan and Guydosh, 2021; Simms *et al*., 2017; Sinha et al., 2020; Sundaramoorthy *et al*., 2017). Paradoxically, our data suggests that ribosome collisions can also do the opposite: suppress recycling of ribosomes. We show that ribosomes translating poly(A) stretches on their own are rapidly removed from the RNA by quality control pathways, but that this ribosome removal is potently suppressed by ribosome cooperativity. Since ribosome cooperativity suppresses ribosome pausing and pausing of single ribosomes results in ribosome removal, it is likely that ribosome cooperativity enhances processivity by suppressing pausing and thus by preventing quality control mechanisms from removing the ribosomes from the mRNA.

Our results reveal for the first time in human cells that single paused ribosomes are efficiently targeted by surveillance pathways. A recent study in yeast identified the E3 ubiquitin ligases Mag2 and Fap1, acting in concert with the RQT ribosome splitting complex, in sensing and clearing of single, decoding-defective ribosome mutants arrested at the start codon (Li *et al*., 2022). Possibly, an analogous quality control pathway exists in human cells, which also relies on the ASC-1 complex and that recognizes and removes single ribosomes paused within the CDS. One key difference between our study and the earlier yeast study (Li *et al*., 2022)-is that in the earlier work, collisions between scanning 40S ribosomal subunits and the 80S ribosome on the start codon may still occur, while in our experiments there is only a single ribosome per RNA, thus unequivocally demonstrating that single paused ribosomes can be targeted by surveillance mechanisms. Removal of single paused ribosomes likely also depends on the state in which the ribosomes is paused, since we find that single ribosomes translating poly(A) sequences are recycled far more efficiently than single ribosome translating Xbp1 or RNA pseudoknot structures, even when pause durations are similar. socRNAs represent an invaluable tool to further dissect quality control pathways targeting single paused ribosomes in the future.

Previous work has also suggested that ribosome collisions induce frameshifting (Champagne et al., 2022; Simms et al., 2019). In the socRNAs used in this study stop codons are present in both alternative reading frames, so events that induce frameshifting would result in abortive translation and thus a reduction in ribosome processivity. Paradoxically, in our experiments processivity is enhanced in the presence of a second ribosome when collisions are frequent. Possibly, frameshifting only occurs in the case of prolonged collision, if collisions cannot resolve the pause. Alternatively, the increased processivity in our assay is caused by a severe suppression of ribosome recycling, masking a parallel, smaller increase in frameshifting that reduces processivity. Directly assessing frameshifting upon collisions of different durations using dual-color socRNAs (Madern et al., 2024) will be an interesting future avenue of investigation.

In summary, we propose that transient collisions can prevent ribosome recycling by suppressing pausing. However, if collisions persist, collisions eventually induce ribosome recycling to clear the mRNA and allow continued translation by other ribosomes. This paradoxical activity of ribosome collisions minimizes waste by limiting unnecessary recycling of ribosomes, while simultaneously preventing prolonged obstruction of mRNAs by permanently stalled ribosomes.

## Acknowledgements

We thank members of the Tanenbaum for helpful discussions. We also thank Jet Segeren for help with experiments and Olivia Rissland, Allen Buskirk, Rachel Green and Marco Catipovic for critical reading of the manuscript. M.F.M., S.Y., and M.E.T. were supported by the Oncode Institute, which is partly funded by the Dutch Cancer Society (KWF). M.E.T. acknowledges funding from the VIDI (NWO/016.VIDI.189.005). S.Y. acknowledges funding from the European Union’s Horizon 2020 research and innovation program under the Marie Skłodowska-Curie grant agreement no. 101026470.

## METHODS

### Cell lines

Human U2OS cells used for socRNA imaging stably express STAb-GFP, ALFANb-CAAX, and tetR. All cells were grown in DMEM (4.5 g/L glucose, Gibco) supplemented with 5% fetal bovine serum (Sigma-Aldrich) and 1% penicillin/streptomycin (Gibco) with 5% CO_2_ at 37°C. Cells were confirmed to be mycoplasma negative.

### Plasmids

The sequences of plasmids used in this study can be found in Table S2.

### Live-cell microscopy

#### Microscopes

Imaging experiments were performed using a Nikon TI inverted microscope with NIS Element Software equipped with a perfect focus system, a Yokagawa CSU-X1 spinning disc, an iXon Ultra 897 EM-CCD camera (Andor), and a motorized piezo stage (Nanocan SP400, Prior). The microscope was equipped with a temperature-controlled box. A 100x 1.49 NA oil-immersion objective was used for all imaging experiments.

#### Cell culture for imaging

Unless noted otherwise, socRNA imaging was performed by seeding cells stably expressing STAb-GFP, ALFANb-CAAX, and TetR in a 96-well glass-botom plate (Matriplates, Brooks Life Science Systems) at ∼25% confluency. The next day, the cells were transfected using Fugene (Promega) with a plasmid encoding the socRNAs of interest. Imaging was done the following day by replacing the medium with pre-warmed imaging medium (CO_2_-independent Leibovitz’s-15 medium (Gibco) containing 5% fetal bovine serum (Sigma-Aldrich) and 1% penicillin/streptomycin (Gibco)). 90 minutes prior to the start of imaging, doxycycline (Dox, 1 µg/mL) was added to the cells to induce socRNA expression. All live-cell imaging experiments were performed at 37°C.

#### Drug treatment

To precisely determine the number of translating ribosomes on socRNAs, the translation inhibitor puromycin (0.1 mg/mL; ThermoFischer Scientific) was added to cells 1-4 hours after the start of imaging to induce nascent chain release. To prevent potential degradation of nascent polypeptides, the proteasome inhibitor MG132 (10 μM) was added at the start of imaging for various live-cell imaging experiments (Figures 2K-2L and 5C-5D). To prevent overloading of the ribosome collision surveillance pathway, harringtonine (3 µg/mL; Cayman Chemical) was added to cells at the start of imaging (Figures 2K-2L).

#### Single-molecule imaging of socRNAs

For live-cell imaging of socRNAs, the x, y positions for imaging were chosen based on the presence of translated socRNAs in cells. Images were acquired every 90-180 sec for 1-4 hours with exposure times for the 488 laser ranging from 50-100 ms. Unless stated otherwise, single z-plane images were acquired with focus on SunTag-GFP foci on the plasma membrane. For experiments in which the GFP fluorescence intensity of individual 24xSunTag arrays was measured, the cells were transfected with a plasmid encoding the 24xSunTag-CAAX protein.

### Ribosome processivity under low doses of elongation inhibitors

To determine the effect of low doses of elongation inhibitors on socRNAs translated by either one or two ribosomes, socRNA expression was induced and cells were selected for imaging. Upon start of imaging, harringtonine was added to cells (3 μg/mL, Cayman Chemical) to induce run-off of all translating ribosomes to prevent the occurrence of widespread ribosome collisions on endogenous mRNAs and potential overloading of collision sensing pathways. 30 minutes after Harringtonine addition, low doses of elongation inhibitors (emetine at 0.01μg/mL, anisomycin at 0.03μg/mL and narciclasine at 0.03μg/mL) was added to cells. Together with the respective elongation inhibitor, MG132 was added to cells to prevent potential degradation of socRNA translation products as a consequence of RQC. socRNA translation sites were imaged and tracked for another 100 minutes, after which puromycin was added to cells to determine the number of translating ribosomes per socRNA. In the case of nascent chain release or ribosome recycling, which results in splitting of the socRNA translation site initially translated by two ribosomes, the intensities of both daughter spots were still followed and measured for inclusion in the analysis.

### siRNA transfections

Cells were first reverse transfected with siRNAs at a final concentration of 10 nM using Lipofectamine RNAiMAX (Invitrogen) and seeded in 96-well glass-botom imaging plates. After 24hr, the cells were trypsinized, transfected with a second dose of 10 nM siRNA, and re-plated in 96-well glass-botom imaging plates. 24 hr after the second siRNA transfection, cells were transfected with plasmid DNA encoding a socRNA, as described above. 24 hr after DNA transfection, imaging experiments or/and qPCR were performed. The sequences of the siRNAs used in this study are listed below.

ZNF598: 5’-ACGAGGAGGUGGACAGGUAUU-3’ (Dharmacon)

ASCC3: 5’-GAUAAAGCGAUCUAAACUUUU-3’ (Dharmacon)

Kif18b (used as a Control siRNA): TTGATGACTGTGGCTGGGC (Dharmacon)

#### siRNA knockdown efficiency

To determine the knock-down efficiency of siRNAs, RNA from siRNA-treated cells was isolated using TRIsure (Bioline). Next, cDNA was synthesized using Random hexamers and Tetro Reverse Transcriptase (Bioline). Quantitative PCRs (qPCRs) were performed using SYBR-Green Supermix (Bio-Rad) on a Bio-Rad Real-time PCR machine (CFX Connect Real-Time PCR Detection System). mRNA levels were determined by qPCR. If the quantitation cycle (C_q_) of a sample was higher than that of a water control, the sample was excluded from analysis. GAPDH and Ribophorin were used as reference genes and fold changes were calculated using the ΔΔCt method. The sequences of the oligonucleotides used for qPCR are listed below (5’ – 3’).

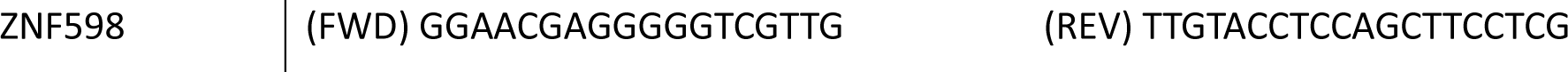
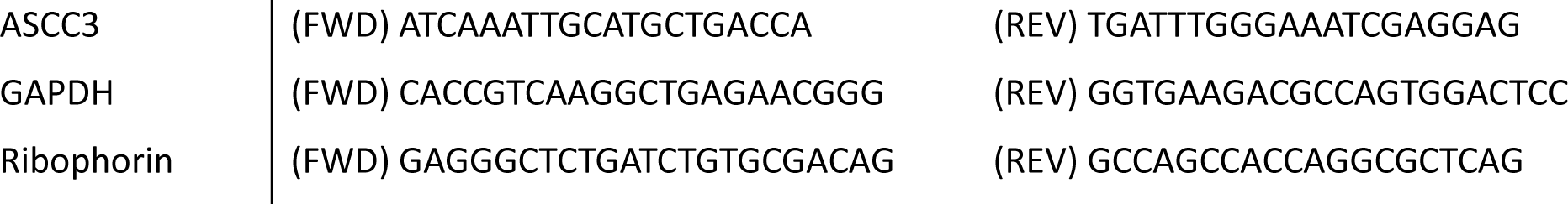

## QUANTIFICATION AND STATISTICAL ANALYSIS

### Post-acquisition processing of microscopy data

For all images, flat-field correction was performed using images obtained from concentrated dye solutions (4 µg/mL DyLight^Tm^ 488 NHS Ester for 488 laser line) and dark current images.

### Tracking and intensity measurements of socRNA foci

For tracking and fluorescence intensity measurements of socRNAs, we used the ‘TransTrack’ software package as previously described (Boersma et al., 2019). All resulting traces underwent manual curation to ensure accuracy.

To correct for photobleaching of membrane-tethered GFP-foci, we used GFP intensity time traces from foci exhibiting no increase in intensity over time, which were acquired in the same imaging experiments. The decrease in fluorescence intensity of these GFP foci over time was fit with a single exponential decay function to determine the bleaching rate. All GFP foci intensities were then corrected for photobleaching. We found that bleach correction on GFP foci rather than whole cell fluorescence is essential, as GFP foci bleach faster than the whole cell, because only a small region of the cell in the z-direction is excited and GFP foci stay within the excitation focus plane throughout the experiment.

### Translation elongation rates of single ribosomes on socRNAs

To determine the number of translating ribosomes per socRNA, puromycin (0.1 mg/mL) was added to cells at the end of the imaging experiment and the number of ribosomes was determined by counting the number of splitting GFP foci after puromycin addition (Figures 1B and 1C). The elongation rate of ribosomes on socRNAs was determined by fitting a linear function to GFP intensity time traces to extract the slope of intensity increase phase before puromycin addition. Also, when GFP foci split into multiple foci over the course of the experiment, we took the intensity traces until the moment of foci splitting to extract the slope of intensity increase phase before foci splitting. The slope was then divided by the number of ribosomes to determine the translation elongation speed per ribosome. To convert rates of GFP intensity increase into the unit of amino acids translated per second, we first determined the intensity of a single GFP molecule under our experimental settings. To achieve this, we measured the intensity of individual ‘mature’ SunTag proteins containing 24 repeats of the SunTag peptide fused to a CAAX motif (24xSunTag-CAAX) using the same settings as those used in the imaging experiment. We divided the average intensity of 24xSunTag-CAAX foci by 24 to obtain the intensity of a single GFP molecule. Using the intensity of a single GFP molecule, we could calculate the number of SunTag epitopes synthesized per unit of time for translated socRNAs. Next, for each socRNA, we calculated the average number of codons that need to be translated for the synthesis of one SunTag epitope; we determined the number of codons for the translation of a full cycle for each socRNA, and the number of SunTag epitopes synthesized upon translation of the socRNA once (equal to the number of SunTags encoded in a socRNA, 5 or 10, unless noted otherwise). Based on the number of codons in one full cycle of socRNA translation and the number of SunTags encoded in a socRNAs, we calculated the elongation rate in amino acids per second.

### Calculating ribosome pause time

To determine ribosome pause time on socRNAs encoding a pause sequence, we determined the average elongation rates (i.e. the total time to complete translation of one full circle, which represents the time needed to translate the non-pause sequence plus the pause time on the pause sequence) of single ribosomes as described in the paragraph above. We then subtracted the average translation time for one cycle of translation of a matched socRNA lacking the pause sequence to obtain the pause duration per cycle.

### Calculating duration of emetine-induced ribosome collisions

We wanted to quantitatively assess the moment from ribosome collision until ribosome recycling in emetine-induced ribosome collisions. GFP foci were tracked before and after emetine addition and the foci intensities were ploted over time. Upon binding of emetine to one of the two ribosomes, a transition from a positive slope to a plateau is observed in the GFP intensity time trace, where the transition point represents the moment of emetine-induced ribosome pausing. To identify this transition time point, we fit the GFP intensity time traces with two distinct linear regression models such that the two state regression model best fit the data using the least squares approach. Importantly, the second of the two linear regression models was constrained to have a slope of zero to fit the paused state, and the first linear regression model needed to exhibit a positive slope to fit the trace up until the moment of ribosome pausing. The intersection point between the two regression models was used to calculate the moment in time of collision onset, and the time interval between collision onset and visible splitting of socRNA translation spot into two daughter spots was calculated. To precisely determine the collision duration from the moment of collision onset to the moment of ribosome recycling, we corrected for the time needed from the moment of ribosome recycling until the two GFP foci have diffused sufficiently far apart from each other to be scored as a ribosome splitting. To determine this time, socRNAs translated by two ribosomes were imaged and treated with puromycin and the time from puro addition until splitting was recorded (Figure S2E, single exponential function: half life = 1.4 min, tau = 2.0). In addition, we also corrected for the average duration needed for two ribosomes to collide upon emetine binding to one of the two ribosomes. For this calculation we assumed that the relative distance between the two ribosomes on the socRNA is half the length of the socRNA (i.e. that they are randomly positioned relative to each other) (average time = 0.50 min on socRNA with a size of 525 nt and elongation speed of 2.6 aa/s). Both the average time needed for GFP foci diffusion and for collision upon emetine binding were subtracted from our data to calculate the duration of ribosome collision until the moment of recycling.

### Calculating the ribosome cooperativity index

The cooperativity index represents a measure for the degree to which the presence of a second ribosome speeds up the translation elongation speed of another ribosome. The cooperativity index was calculated for socRNAs containing a pause site. To calculate the cooperativity index, we first calculated the average total pause time per full cycle of socRNA translation per ribosome. The total pause time includes both the pause on the pause site (e.g. Xbp1 sequence) and the ‘waiting time’ in which a trailing ribosome is paused upstream of a ribosome that is paused on the pause site (which is referred to as ribosome interference). The total pause time is calculated from the ribosome interference simulation as described in *Ribosome interference during translation on socRNAs with pause sequence* (e.g., black data in Figures 4A and 4B). To calculate the cooperativity index, the calculated total pause time per ribosome was divided by the pause time per ribosome determined from the experiments (e.g., cyan data in Figures 4A and 4B). The cooperativity index indicates how much the total pause time is reduced by second ribosome.

### Identifying transient pauses in GFP intensity time traces from translated socRNAs

To identify pauses within the intensity time traces (Figures 6A and 6B), the intensity value of each time point in the raw intensity traces (black line in Figure 6A) was first divided by the number of ribosomes on that socRNA to calculate the elongation rate per ribosome. Next, the intensity traces were smoothed using a 6-point moving median to eliminate outlier data points. Subsequently, a 5-point moving average was applied to further smooth the data (red line in Figure 6A). Following smoothing, the first derivative, which represents the differences between adjacent intensities, was calculated (black line in Figure 6B). Pause identification was performed using a Hidden Markov Model (vbFRET algorithm (Bronson et al., 2009)) with default settings of the algorithm. To compare pause frequency between one ribosome traces and multiple ribosomes traces, identical threshold for defining pause state were used for both data set.

### Quantification of ribosome processivity

To determine the number of codons translated by individual ribosomes on socRNAs, we tracked GFP intensity time traces for translated socRNAs and determined the moment when the GFP intensity stopped increasing for individual translated socRNAs. For socRNAs translated until puromycin addition, we noted the last frame before puromycin addition as the last time-point in which translation was detected. We then measured for each individual socRNA the GFP foci intensity at the last time-point of translation and calculated the total number of codons translated during the experiment based on this final time-point GFP intensity, as described in *Translation elongation rates of single ribosomes on socRNAs*. The fraction of translated socRNAs remaining was then ploted against the total number of codons translated in Kaplan-Meier survival plots.

For socRNAs that were translated by two ribosomes, we determined the moment when the GFP foci split into two foci and kept tracked the intensity of both foci to determine whether one of foci intensities continued increasing (indicating that one of the two ribosomes continued translating) or whether both foci stop increasing in intensity (indicating that both ribosomes aborted translation) after splitting. We then measured the GFP foci intensity at the moment of foci splitting and calculated the total number of codons translated based on the GFP intensity. Next, the total number of translated codons was divided by two to determine the number of codons translated by each ribosome. To compare processivity of single or multiple ribosomes translating a socRNA, the fraction of translating ribosome was ploted against the total number of translated codons per ribosome between as Kaplan-Meier survival curves. For socRNAs translated by two ribosomes, we determine the moment of splitting (indicative of the first ribosome aborting translation), and noted that moment as aborted translation for one ribosome and as the last moment that translation could be detected for the other ribosomes.

### Statistical analyses and generation of graphs

All graphs were generated using Prism GraphPad (v9) or in python 3.10 using Matplotlib. Details of statistical tests for each graph are explained in figure legends.

## SIMULATION

### Modelling ribosome translocation dynamics *in silico*

To investigate the prevalence of ribosome collisions on mRNAs, we simulated translation of ribosomes over time using a computational model. In the model, ribosomes moved one codon at a time with a variable rate, representing the stochastic behavior of ribosome translocation. The time it takes a ribosome to move one codon was determined by randomly selecting a value from a distribution that represented the elongation cycle duration. This elongation cycle duration distribution was constructed in the following way:

1) We determined that the average of the elongation cycle distribution is 3 aa/s, in accordance with previously calculated elongation speed values on linear mRNAs in our cell line (Yan *et al*., 2016).
2) Based on a previously published CryoEM dataset of ribosome elongation (Behrmann *et al*., 2015), the entire elongation cycle was first divided into 7 distinct sub-steps that ribosomes cycle through.
3) The relative abundance of each individual sub-step structure (0.0833, 0.0315, 0.0833, 0.25, 0.396, 0.0833 and 0.073 determined in (Behrmann *et al*., 2015)) was used to first calculate the average duration of each sub-step. Specifically, average sub-step durations were calculated by multiplying the relative abundance of each sub-step by the duration of the entire elongation cycle. Next, for each sub-step, a duration distribution was constructed following a single exponential decay function with a mean determined as described above (See Figure 1E). The sum of the averages from all 7 sub-step distributions is exactly 3 aa/s, equal to the ribosome elongation speed as previously determined on linear mRNAs (Yan *et al*., 2016).
4) To construct the distribution of durations for one entire elongation cycle, a duration for each individual sub-step was randomly selected from the distributions constructed in 3) and these seven durations were summed. This process was repeated 5 million times to construct the distribution of durations of one elongation cycle (See Figure 1F).
5) For each translated codon, we randomly picked a value from the distribution described in 4) to determine how long it takes to translate that codon.
6) To simulate the experimentally observed heterogeneity in elongation rates for individual ribosomes (see Madern et al., 2024 for intrinsic ribosome elongation rate heterogeneity), each individual ribosome was assigned a relative elongation speed picked at random from a Gaussian distribution which was based on the actual elongation speed heterogeneity (mean of 1, coefficient of variation = 0.158). For every individual ribosome, each new elongation cycle duration was divided by the relative elongation speed of that ribosome.

In all simulations, the ribosome footprint size was set to exactly 10 codons, and ribosomes colliding with a slower leading ribosome could not overtake the leading ribosome.

For simulations in which the elongation cycle consisted of 1 rate-limiting sub-step (See Figures S1C, and S1E-S1H), we exchanged the values described in 2) such that one sub-step took up two thirds of the entire elongation cycle duration. Specifically, the 7 values were changed to 0.667, 0.056, 0.056, 0.056, 0.056, 0.056 and 0.056.

### Simulating ribosome collisions on endogenous linear mRNAs (Figure 1)

Translation initiation was simulated using an average interval of 30 seconds between initiation events, in accordance with experimentally determined ribosome initiation rates (Livingston *et al*., 2023; Yan *et al*., 2016). To stimulate the stochasticity of translation initiation kinetics, the interval between two initiation events was picked randomly from a distribution that followed a single exponential decay with a mean of 30 s. Since a ribosome cannot initiate translation until the previous ribosome is at least 10 codons downstream of the initiation site, we introduced a minimum interval between initiation events of 10 s, reflecting the combined time of translation initiation and translation of 10 codons. After initiation, ribosome translocation was simulated for 30 min, as described above. For each simulated mRNA, the fraction of all ribosomes which at some point collided with another ribosome was calculated.

### Simulating ribosome collisions on socRNAs (Figure 1)

To model the time and number of codons translated (per ribosomes) until the first collision between 2 ribosomes translating the same socRNA, ribosomes were initially loaded at random but nonoverlapping locations on the socRNA. To match our experimental data, the socRNA used in this simulation had a length of 175 codons and ribosomes translated with an average speed of 2.6 aa/s, consistent with our experimental measurements. In addition, intrinsic ribosome heterogeneity in speed was incorporated in this simulation, as described above. Translation elongation was initiated *in silico* and the time and number of codons translated until the first collision occurred were recorded for 1000 simulations.

### Ribosome interference during translation on socRNAs without pause sequence

To quantitatively assess how translating ribosomes influence each others translocation dynamics on socRNAs, we conducted simulations in which two ribosomes translate the same socRNA simultaneously.

We used the following parameters for these simulations:

(1) We used the same socRNA lengths as those used in the experiments (in number of codons).
(2) We randomly selected a value for the intrinsic, average ribosome speed for each ribosome, based on the intrinsic ribosome speed distribution. This distribution was obtained from the same cell line (mean = average ribosome elongation speed (aa/sec) acquired from experimental data, coefficient of variation = 0.16) (Madern et al., 2024).
(4) We determined the translation initiation time for each ribosome as described in *Modelling ribosome translocation dynamics in silico*
(5) A ribosome can only move forward by one codon if the second ribosome is >10 codons away (based on a ribosome footprint of 30 nt = 10 codons).

Using the parameter described above, combined with the translation elongation dynamics simulations described in *Modelling ribosome translocation dynamics in silico,* we conducted simulations of ribosome translocation and measured ribosome translocation rates.

We converted total amino acids translated by each ribosome into a GFP intensity time trace to mimic experimental data (Figures 3A and 3B). To do this, we used the single GFP molecule intensity as described in *Translation elongation rates of single ribosomes on socRNAs.* For example, for the socRNA template encoding 10xSunTag, the total number of codons for one cycle is 321 codons and the total number of GFPs synthesized in one cycle is 10. We thus divided 10x(single GFP intensity) by 321 to determine the average GFP increase per codon.

### Ribosome interference during translation on socRNAs with pause sequence

In the simulations where two ribosomes translate the same socRNA encoding a pause sequence, we used the same computational framework described above, but implemented the following the changes:

(1) A pause site was inserted into the socRNA.
(2) A ribosome undergoes a pause when it translates the last codon of the pause site.
(3) The pause time for each pausing event is determined by randomly drawing a value from an exponential decay distribution with a mean value that is based on experimental measurements for socRNAs containing that pause sequence. For mean pause durations the pause time for single ribosomes translating the socRNA was used.

### Slipstream effect with RNA structures

In simulations where two ribosomes translate the same socRNA encoding a pseudoknot RNA structure, we introduced the following adjustments to account for the ‘slipstream’ effect associated with translation of an RNA structure by multiple ribosomes:

(1) A pseudoknot, spanning a length of 26 codons, was inserted into the socRNA.
(2) To simulate ribosome pausing upstream of the pseudoknot, we incorporated the requirement in our simulations that when a ribosome has codon ‘i’ in the A site, translocation to the next codon requires the unwinding of codon ‘i+4’ downstream (Qu *et al*., 2011). This results in a ribosome pause occurring 4 codons upstream of the first codon of the pseudoknot.
(3) The duration of ribosome pausing upstream of the pseudoknot is equal to the unfolding time of the pseudoknot structure, which was determined in our simulations by randomly drawing a value from an exponential decay distribution with a mean value equal to the average pause time of single ribosomes on socRNAs encoding the pseudoknot.
(4) After unfolding of the pseudoknot structure, the ribosome resumes translation and translates the pseudoknot sequence at the same speed as non-pseudoknot sequences are translated at.
(5) Slipstream effect: if a ribosome translates the codon that is 4 codons upstream of the first codon of the pseudoknot (referred to as ‘i’), and the other ribosome (which has a 30 nt footprint) is still covering part of the pseudoknot sequence with its footprint, the trailing ribosome does not pause at codon ‘i-4’. This is because the leading ribosome has already unfolded the pseudoknot structure and prevents its refolding when the ribosome footprint covers the pseudoknot sequence.

### Ribosome cooperativity on socRNAs with a pause sequence

To incorporate ribosome cooperativity in the simulation where two ribosomes translate the same socRNA encoding a pause sequence, the same parameters were used as described in *Ribosome interference during translation on socRNAs with pause sequence*, but with the following modifications:

When a leading ribosome pauses at the pause site (codon ‘i’) and a trailing ribosome has codon ‘i-5’, in the A site, i.e., a translating ribosome collides with the paused ribosome, the paused ribosome automatically translocates to the next codon, irrespective of the pause duration that was drawn from the pause time distribution. In the simulations in figures S3A and S3B, we also included a delay time between collision and automatic translocation of the paused ribosome. We used different delays after collision (e.g., 0, 10, 20, 40 seconds for Xbp1(S255A)). In the simulations in figures S3C and S3D, we included different probabilities of cooperativity for which collision-induced resumption of translation happens with certain probability. For example, 0 % indicates no cooperativity as same as ribosome interference simulation.

---

**Figure S1.**
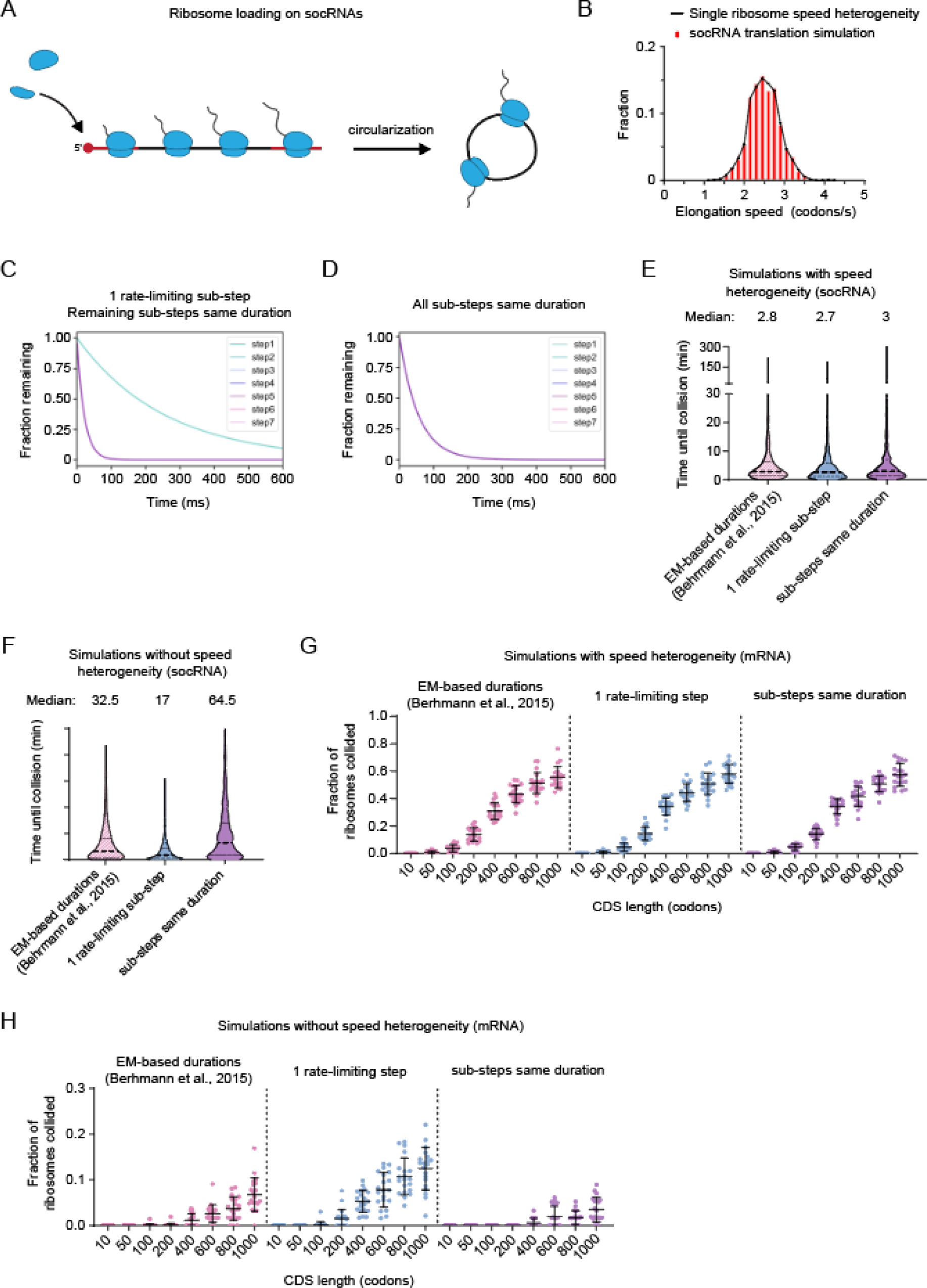
related to Figure 1. Simulating ribosome collisions on socRNAs and linear mRNAs. A) Schematic depicting the mechanism by which ribosomes are loaded onto socRNAs. Ribosomes are first recruited to the 5’end of the linear precursor RNA through 5’ cap-dependent recruitment. While ribosomes are translating the coding sequence of the linear pre-cursor RNA, the internal section of the linear RNA becomes circularized (black), essentially trapping ribosomes on the socRNA. B) The distribution of single ribosome elongation speeds determined experimentally (black line) (Madern et al., 2024) or from the simulations (red bars, see Methods). C-D) Distributions of the duration of each step in the translation elongation cycle used in simulations. E-F) The time until the first collision between two ribosomes translating the same socRNA was determined based on simulations. Simulations were repeated 1000 times for each condition (See Figure 1E and S1C-D). Thick dashed line indicates the median, thin lines indicate 25th and 75th percentile. Median values are listed above each graph. E) Simulations including ribosome speed heterogeneity were performed for different elongation cycle sub-step regimes to determine the time until 2 ribosomes translating the same socRNA collide. Data showing ‘EM-based durations’ are reploted from Figure 1H. F) Simulations without including ribosome speed heterogeneity were performed for different elongation cycle sub-step regimes to determine the time until 2 ribosomes translating the same socRNA collide. G-H) Simulations were performed to determine the frequency of ribosome collisions on mRNA molecules as a function of mRNA coding sequence (CDS) length. Each data point represents a single simulated mRNA molecule (See Methods), and 30 mRNAs were simulated for each different condition (See Figure 1E and S1C-D). Error bars represent standard deviation. G) Simulations of mRNA translation including intrinsic ribosome speed heterogeneity were performed to determine the fraction of ribosomes undergoing at least one collision during translation of the CDS. H) Simulations of mRNA translation without intrinsic ribosome speed heterogeneity were performed to determine the fraction of ribosomes undergoing at least one collision during translation of the CDS. Error bars indicate standard deviation from independent experiments.

**Figure S2.**
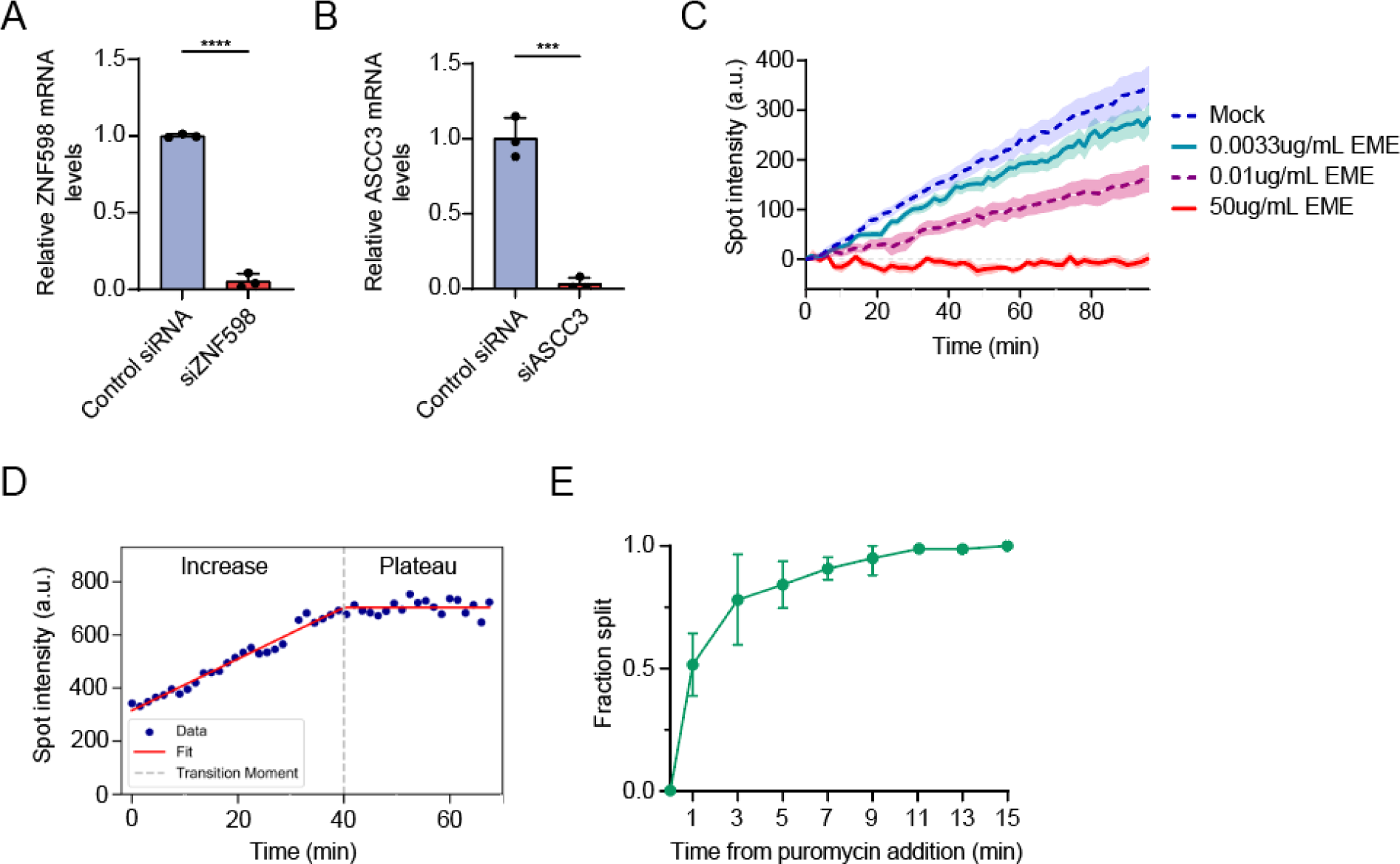
related to Figure 2. Controls for investigating ribosome collisions and recycling. A-B) Quantitative PCRs were performed to assess the knockdown efficiency of siRNA treatment for ZNF598 (B) and ASCC3 (C). ***, **** indicate p<0.001, and 0.0001, respectively, determined by t-test. Data represent mean ± SD of independent experiments. Dots represent the data from independent experiments. C) U2OS cells stably expressing STAb-GFP, ALFANb-CAAX, and TetR were transfected with control socRNAs and imaged by time-lapse microscopy. Emetine was added at different concentrations to determine the dose-dependent effect on average protein synthesis rates by single translating ribosomes. Dashed lines represent data reploted from Figures 3I-3J. Lines indicate mean values and shaded regions indicate 95% CI. D) Representative example of data fitting approach to identify the plateau duration preceding the moment of ribosome recycling for trace shown in Figure 2K. E) U2OS cells stably expressing STAb-GFP, ALFANb-CAAX, and TetR were transfected with control socRNAs and imaged by time-lapse microscopy. Puromycin was added to cells to determine the time needed for nascent chains from socRNAs translated by two ribosomes to visibly dissociate from each other. Fitting with a single exponential function reveals a half life of 1.4 minutes. The number of experimental repeats and cells analyzed per experiment are listed in Table S1.

**Figure S3.**
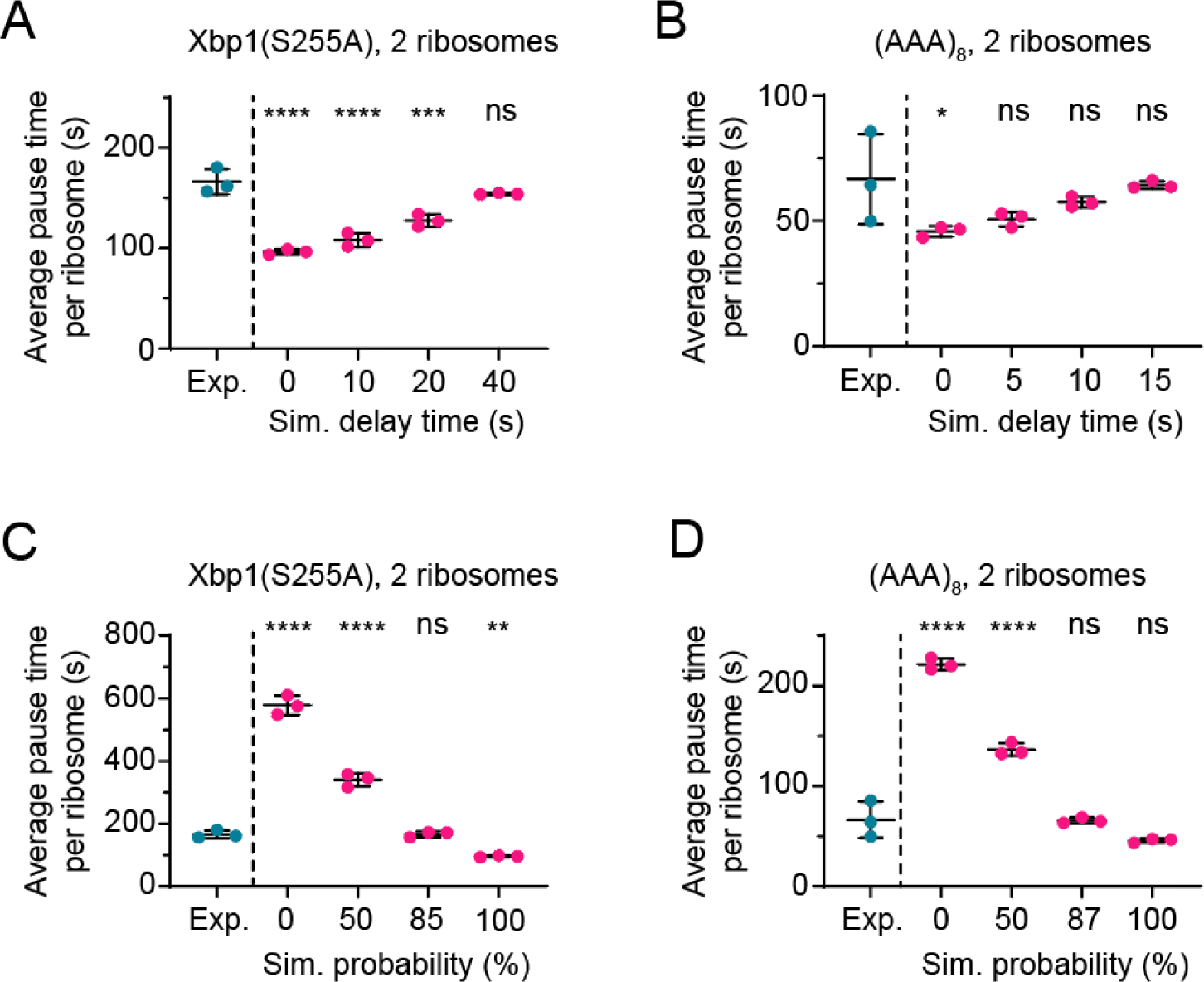
related to Figure 4. Simulation of ribosome cooperativity. A-D) Simulation of the average pause duration on Xbp1(S255A) (A,C) or (AAA)_8_ (B,D) pause sequences for socRNAs translated by two ribosomes. Collision-induced resumption of translation of the paused ribosome was simulated for two different ribosome cooperativity models. Cyan (experimental data) dots are reploted from figure 3D for comparison. A-B) Different delay times between the moment of collision and resumption of translation of the paused ribosome were simulated. C-D) For each collision between a translocating ribosome and a ribosome paused on indicated pause sequences a probability was simulated that the collision resulted in resumption of translation. A probability of 0% means that no ribosome cooperativity was included. *, **, ***, **** indicate p<0.05, 0.01, 0.001, and 0.0001, respectively, determined by t-test. The horizontal black lines and error bars represent mean ± SD of independent experiments or simulations. The dots represent the data from independent experiments or simulations.

**Figure S4.**
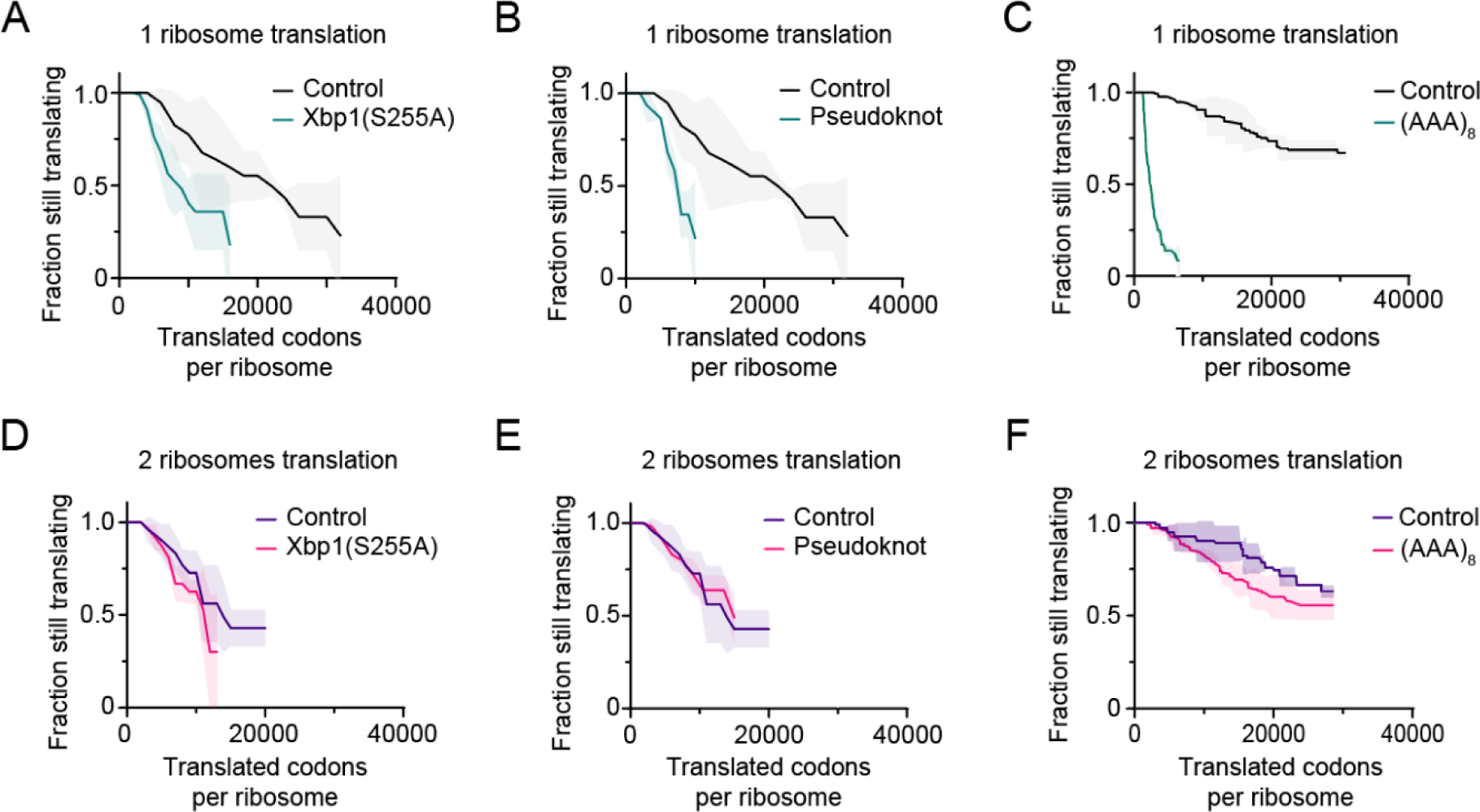
related to Figure 5. Ribosome cooperativity enhances processivity. A-F) U2OS cells stably expressing STAb-GFP, ALFANb-CAAX, and TetR were transfected with indicated socRNAs and imaged by time-lapse microscopy. Kaplan-Meyer survival curve of indicated socRNAs shows the total number of codons translated by individual ribosomes before aborting translation. Xbp1(S255A) (A, D),Pseudoknot (B,E) and (AAA)_8_ (C,F) data is reploted from Figure 5A for comparison. Data from the control socRNAs (socRNAs lacking a pause sequence) (C, F) is reploted from Figure 1J for comparison. Lines indicate mean values and shaded regions indicate standard deviation. The number of experimental repeats and cells analyzed per experiment are listed in Table S1.

**Video S1 – related to Figure 1. Long-term imaging of ribosomes.**

U2OS cells stably expressing STAb-GFP, ALFANb-CAAX, and TetR were transfected with a plasmid encoding a socRNA with 5xSunTag and 1xALFA-tag and imaged by time-lapse microscopy. Images were acquired every 2 min. Puromycin was added at minute 192 to determine the number of ribosomes translating on each socRNA. Scale bar, 10 μm.

**Video S2 – related to Figure 1. Release of nascent chains by puromycin enables quantification of the number of ribosomes translating each socRNA.**

U2OS cells stably expressing STAb-GFP, ALFANb-CAAX, and tetR were transfected with a plasmid encoding a socRNA with 10xSunTag and 2xALFA-tag. Puromycin was added to socRNA-expressing cells at t=27 min to induce nascent chain release and to allow scoring of the number of ribosomes per socRNA. Images were acquired every 90 seconds. Scale bar, 1 μm.

**Video S3 – related to Figure 1. Animation of a simulation of linear mRNA translation by ribosomes with elongation speed heterogeneity.**

Representative movie of a simulation used to determine the frequency of ribosome collisions on linear mRNAs (see Figure 1M). Experimentally determined values for ribosome elongation speed heterogeneity were included in the simulation (see Methods). In the movie, the position of each simulated ribosome is shown after every elapsed second, and translating ribosomes light up red if they have collided with a trailing ribosome within the last second.

**Video S4 – related to Figure 1. Animation of a simulation of linear mRNA translation by ribosomes without elongation speed heterogeneity.**

Representative movie of a simulation used to determine the frequency of ribosome collisions on linear mRNAs (see Figure S1H, EM-based durations). Experimentally determined values for ribosome elongation speed heterogeneity were not included in this simulation. In the movie, the position of each simulated ribosome is shown after every elapsed second, and translating ribosomes light up red if they have collided with a trailing ribosome within the last second.

**Video S5 – related to Figure 3. Animation of a simulation of ribosome interference on socRNAs containing a pause site.**

Simulation of ribosome interference during translation on socRNAs with a pause site. Two ribosomes translate a socRNA of 326 codons at an elongation rate of 2.5 codons/s. Pause time at the pause site follows an exponential distribution with mean 340 seconds based on experimentally determined values (See Methods). Ribosome interference becomes more pronounced when a paused ribosome at the pause site blocks translocation of the trailing ribosome.

**Video S6 – related to Figure 4. Animation of a simulation of ribosome cooperativity on socRNAs containing a pause site.**

Simulation of ribosome cooperativity during translation on a socRNAs with a pause site. Two ribosomes translate a socRNA of 326 codons at an elongation rate of 2.5 codons/s. Pause time at the pause site follows an exponential distribution with mean 340 seconds determined from Xbp1 socRNA experiments. In this simulation ribosome collisions instantaneously resolve the pause of the leading ribosome.

## References

Arthur, L., Pavlovic-Djuranovic, S., Smith-Koutmou, K., Green, R., Szczesny, P., and Djuranovic, S. (2015). Translational control by lysine-encoding A-rich sequences. Sci Adv 1. 10.1126/sciadv.1500154.

Behrmann, E., Loerke, J., Budkevich, T.V., Yamamoto, K., Schmidt, A., Penczek, P.A., Vos, M.R., Burger, J., Mielke, T., Scheerer, P., and Spahn, C.M. (2015). Structural snapshots of actively translating human ribosomes. Cell 161, 845–857. 10.1016/j.cell.2015.03.052.

Caliskan, N., Katunin, V.I., Belardinelli, R., Peske, F., and Rodnina, M.V. (2014). Programmed -1 frameshifting by kinetic partitioning during impeded translocation. Cell 157, 1619–1631. 10.1016/j.cell.2014.04.041.

Champagne, J., Mordente, K., Nagel, R., and Agami, R. (2022). Slippy-Sloppy translation: a tale of programmed and induced-ribosomal frameshifting. Trends Genet 38, 1123–1133. 10.1016/j.tig.2022.05.009.

Chandrasekaran, V., Juszkiewicz, S., Choi, J., Puglisi, J.D., Brown, A., Shao, S., Ramakrishnan, V., and Hegde, R.S. (2019). Mechanism of ribosome stalling during translation of a poly(A) tail. Nat Struct Mol Biol 26, 1132–1140. 10.1038/s41594-019-0331-x.

Chen, C.-W., and Tanaka, M. (2018). Genome-wide Translation Profiling by Ribosome-Bound tRNA Capture. Cell Reports 23, 608–621. 10.1016/j.celrep.2018.03.035.

Collart, M.A., and Weiss, B. (2020). Ribosome pausing, a dangerous necessity for co-translational events. Nucleic Acids Res 48, 1043–1055. 10.1093/nar/gkz763.

D’Orazio, K.N., and Green, R. (2021). Ribosome states signal RNA quality control. Mol Cell 81, 1372–1383. 10.1016/j.molcel.2021.02.022.

Dever, T.E., Dinman, J.D., and Green, R. (2018). Translation Elongation and Recoding in Eukaryotes. Cold Spring Harb Perspect Biol 10. 10.1101/cshperspect.a032649.

Dmitriev, S.E., Vladimirov, D.O., and Lashkevich, K.A. (2020). A Quick Guide to Small-Molecule Inhibitors of Eukaryotic Protein Synthesis. Biochemistry (Mosc) 85, 1389–1421. 10.1134/S0006297920110097.

Eiler, D.R., Wimberly, B.T., Bilodeau, D.Y., Taliaferro, J.M., Reigan, P., Rissland, O.S., and Kieft, J.S. (2024). The Giardia lamblia ribosome structure reveals divergence in several biological pathways and the mode of emetine function. Structure. 10.1016/j.str.2023.12.015.

Ferguson, A., Wang, L., Altman, R.B., Terry, D.S., Juete, M.F., Burnett, B.J., Alejo, J.L., Dass, R.A., Parks, M.M., Vincent, C.T., and Blanchard, S.C. (2015). Functional Dynamics within the Human Ribosome Regulate the Rate of Active Protein Synthesis. Mol Cell 60, 475–486. 10.1016/j.molcel.2015.09.013.

Garreau de Loubresse, N., Prokhorova, I., Holtkamp, W., Rodnina, M.V., Yusupova, G., and Yusupov, M. (2014). Structural basis for the inhibition of the eukaryotic ribosome. Nature 513, 517–522. 10.1038/nature13737.

Garzia, A., Jafarnejad, S.M., Meyer, C., Chapat, C., Gogakos, T., Morozov, P., Amiri, M., Shapiro, M., Molina, H., Tuschl, T., and Sonenberg, N. (2017). The E3 ubiquitin ligase and RNA-binding protein ZNF598 orchestrates ribosome quality control of premature polyadenylated mRNAs. Nat Commun 8, 16056. 10.1038/ncomms16056.

Gobet, C., Weger, B.D., Marquis, J., Martin, E., Neelagandan, N., Gachon, F., and Naef, F. (2020). Robust landscapes of ribosome dwell times and aminoacyl-tRNAs in response to nutrient stress in liver. Proceedings of the National Academy of Sciences 117, 9630–9641. doi:10.1073/pnas.1918145117.

Goldman, D.H., Livingston, N.M., Movsik, J., Wu, B., and Green, R. (2021). Live-cell imaging reveals kinetic determinants of quality control triggered by ribosome stalling. Mol Cell 81, 1830–1840 e1838. 10.1016/j.molcel.2021.01.029.

Gutierrez, E., Shin, B.S., Woolstenhulme, C.J., Kim, J.R., Saini, P., Buskirk, A.R., and Dever, T.E. (2013). eIF5A promotes translation of polyproline motifs. Mol Cell 51, 35–45. 10.1016/j.molcel.2013.04.021.

Hill, C.H., and Brierley, I. (2023). Structural and Functional Insights into Viral Programmed Ribosomal Frameshifting. Annu Rev Virol 10, 217–242. 10.1146/annurev-virology-111821-120646.

Hoek, T.A., Khuperkar, D., Lindeboom, R.G.H., Sonneveld, S., Verhagen, B.M.P., Boersma, S., Vermeulen, M., and Tanenbaum, M.E. (2019). Single-Molecule Imaging Uncovers Rules Governing Nonsense-Mediated mRNA Decay. Mol Cell 75, 324–339 e311. 10.1016/j.molcel.2019.05.008.

Holm, M., Natchiar, S.K., Rundlet, E.J., Myasnikov, A.G., Watson, Z.L., Altman, R.B., Wang, H.Y., Taunton, J., and Blanchard, S.C. (2023). mRNA decoding in human is kinetically and structurally distinct from bacteria. Nature 617, 200–207. 10.1038/s41586-023-05908-w.

Ikeuchi, K., Tesina, P., Matsuo, Y., Sugiyama, T., Cheng, J., Saeki, Y., Tanaka, K., Becker, T., Beckmann, R., and Inada, T. (2019). Collided ribosomes form a unique structural interface to induce Hel2-driven quality control pathways. EMBO J 38. 10.15252/embj.2018100276.

Inada, T. (2017). The Ribosome as a Platform for mRNA and Nascent Polypeptide Quality Control. Trends in Biochemical Sciences 42, 5–15. 10.1016/j.tibs.2016.09.005.

Juszkiewicz, S., Chandrasekaran, V., Lin, Z., Kraatz, S., Ramakrishnan, V., and Hegde, R.S. (2018). ZNF598 Is a Quality Control Sensor of Collided Ribosomes. Mol Cell 72, 469–481 e467. 10.1016/j.molcel.2018.08.037.

Juszkiewicz, S., and Hegde, R.S. (2017). Initiation of Quality Control during Poly(A) Translation Requires Site-Specific Ribosome Ubiquitination. Mol Cell 65, 743–750 e744. 10.1016/j.molcel.2016.11.039.

Juszkiewicz, S., Speldewinde, S.H., Wan, L., Svejstrup, J.Q., and Hegde, R.S. (2020). The ASC-1 Complex Disassembles Collided Ribosomes. Mol Cell 79, 603–614 e608. 10.1016/j.molcel.2020.06.006.

Knight, J.R.P., Garland, G., Poyry, T., Mead, E., Vlahov, N., Sfakianos, A., Grosso, S., De-Lima-Hedayioglu, F., Mallucci, G.R., von der Haar, T., et al. (2020). Control of translation elongation in health and disease. Dis Model Mech 13. 10.1242/dmm.043208.

Li, S., Ikeuchi, K., Kato, M., Buschauer, R., Sugiyama, T., Adachi, S., Kusano, H., Natsume, T., Berninghausen, O., Matsuo, Y., et al. (2022). Sensing of individual stalled 80S ribosomes by Fap1 for nonfunctional rRNA turnover. Mol Cell 82, 3424–3437 e3428. 10.1016/j.molcel.2022.08.018.

Livingston, N.M., Kwon, J., Valera, O., Saba, J.A., Sinha, N.K., Reddy, P., Nelson, B., Wolfe, C., Ha, T., Green, R., et al. (2023). Bursting translation on single mRNAs in live cells. Mol Cell 83, 2276–2289 e2211. 10.1016/j.molcel.2023.05.019.

Madern, M.F., Yang, S., Witeveen, O., Bauer, M., Tanenbaum, M.E. (2024). Dissecting translation elongation dynamics through ultra-long tracking of single ribosomes. bioRxiv

Matsuo, Y., Ikeuchi, K., Saeki, Y., Iwasaki, S., Schmidt, C., Udagawa, T., Sato, F., Tsuchiya, H., Becker, T., Tanaka, K., et al. (2017). Ubiquitination of stalled ribosome triggers ribosome-associated quality control. Nat Commun 8, 159. 10.1038/s41467-017-00188-1.

Matsuo, Y., Tesina, P., Nakajima, S., Mizuno, M., Endo, A., Buschauer, R., Cheng, J., Shounai, O., Ikeuchi, K., Saeki, Y., et al. (2020). RQT complex dissociates ribosomes collided on endogenous RQC substrate SDD1. Nature Structural & Molecular Biology 27, 323–332. 10.1038/s41594-020-0393-9.

Matsuo, Y., Uchihashi, T., and Inada, T. (2023). Decoding of the ubiquitin code for clearance of colliding ribosomes by the RQT complex. Nature Communications 14, 79. 10.1038/s41467-022-35608-4.

Meydan, S., and Guydosh, N.R. (2021). A cellular handbook for collided ribosomes: surveillance pathways and collision types. Curr Genet 67, 19–26. 10.1007/s00294-020-01111-w.

Morisaki, T., Lyon, K., DeLuca, K.F., DeLuca, J.G., English, B.P., Zhang, Z., Lavis, L.D., Grimm, J.B., Viswanathan, S., Looger, L.L., et al. (2016). Real-time quantification of single RNA translation dynamics in living cells. Science 352, 1425–1429. doi:10.1126/science.aaf0899.

Namy, O., Moran, S.J., Stuart, D.I., Gilbert, R.J., and Brierley, I. (2006). A mechanical explanation of RNA pseudoknot function in programmed ribosomal frameshifting. Nature 441, 244–247. 10.1038/nature04735.

Niu, X., Sun, R., Chen, Z., Yao, Y., Zuo, X., Chen, C., and Fang, X. (2021). Pseudoknot length modulates the folding, conformational dynamics, and robustness of Xrn1 resistance of flaviviral xrRNAs. Nature Communications 12, 6417. 10.1038/s41467-021-26616-x.

Pichon, X., Bastide, A., Safieddine, A., Chouaib, R., Samacoits, A., Basyuk, E., Peter, M., Mueller, F., and Bertrand, E. (2016). Visualization of single endogenous polysomes reveals the dynamics of translation in live human cells. Journal of Cell Biology 214, 769–781. 10.1083/jcb.201605024.

Qu, X., Wen, J.D., Lancaster, L., Noller, H.F., Bustamante, C., and Tinoco, I., Jr. (2011). The ribosome uses two active mechanisms to unwind messenger RNA during translation. Nature 475, 118–121. 10.1038/nature10126.

Rodnina, M.V., Fischer, N., Maracci, C., and Stark, H. (2017). Ribosome dynamics during decoding. Philos Trans R Soc Lond B Biol Sci 372. 10.1098/rstb.2016.0182.

Saito, K., Kratzat, H., Campbell, A., Buschauer, R., Burroughs, A.M., Berninghausen, O., Aravind, L., Green, R., Beckmann, R., and Buskirk, A.R. (2022). Ribosome collisions induce mRNA cleavage and ribosome rescue in bacteria. Nature 603, 503–508. 10.1038/s41586-022-04416-7.

Shanmuganathan, V., Schiller, N., Magoulopoulou, A., Cheng, J., Braunger, K., Cymer, F., Berninghausen, O., Beatrix, B., Kohno, K., von Heijne, G., and Beckmann, R. (2019). Structural and mutational analysis of the ribosome-arresting human XBP1u. Elife 8. 10.7554/eLife.46267.

Simms, C.L., Yan, L.L., Qiu, J.K., and Zaher, H.S. (2019). Ribosome Collisions Result in +1 Frameshifting in the Absence of No-Go Decay. Cell Rep 28, 1679–1689 e1674. 10.1016/j.celrep.2019.07.046.

Simms, C.L., Yan, L.L., and Zaher, H.S. (2017). Ribosome Collision Is Critical for Quality Control during No-Go Decay. Mol Cell 68, 361–373 e365. 10.1016/j.molcel.2017.08.019.

Sinha, N.K., Ordureau, A., Best, K., Saba, J.A., Zinshteyn, B., Sundaramoorthy, E., Fulzele, A., Garshott, D.M., Denk, T., Thoms, M., et al. (2020). EDF1 coordinates cellular responses to ribosome collisions. Elife 9. 10.7554/eLife.58828.

Snieckute, G., Ryder, L., Vind, A.C., Wu, Z., Arendrup, F.S., Stoneley, M., Chamois, S., Martinez-Val, A., Leleu, M., Dreos, R., et al. (2023). ROS-induced ribosome impairment underlies ZAKα-mediated metabolic decline in obesity and aging. Science 382, eadf3208. doi:10.1126/science.adf3208.

Stoneley, M., Harvey, R.F., Mulroney, T.E., Mordue, R., Jukes-Jones, R., Cain, K., Lilley, K.S., Sawarkar, R., and Willis, A.E. (2022). Unresolved stalled ribosome complexes restrict cell-cycle progression after genotoxic stress. Mol Cell 82, 1557–1572 e1557. 10.1016/j.molcel.2022.01.019.

Sundaramoorthy, E., Leonard, M., Mak, R., Liao, J., Fulzele, A., and Bennett, E.J. (2017). ZNF598 and RACK1 Regulate Mammalian Ribosome-Associated Quality Control Function by Mediating Regulatory 40S Ribosomal Ubiquitylation. Mol Cell 65, 751–760 e754. 10.1016/j.molcel.2016.12.026.

Tanenbaum, M.E., Gilbert, L.A., Qi, L.S., Weissman, J.S., and Vale, R.D. (2014). A protein-tagging system for signal amplification in gene expression and fluorescence imaging. Cell 159, 635–646. 10.1016/j.cell.2014.09.039.

Tesina, P., Lessen, L.N., Buschauer, R., Cheng, J., Wu, C.C., Berninghausen, O., Buskirk, A.R., Becker, T., Beckmann, R., and Green, R. (2020). Molecular mechanism of translational stalling by inhibitory codon combinations and poly(A) tracts. EMBO J 39, e103365. 10.15252/embj.2019103365.

Wang, C., Han, B., Zhou, R., and Zhuang, X. (2016). Real-Time Imaging of Translation on Single mRNA Transcripts in Live Cells. Cell 165, 990–1001. 10.1016/j.cell.2016.04.040.

Wong, W., Bai, X.C., Brown, A., Fernandez, I.S., Hanssen, E., Condron, M., Tan, Y.H., Baum, J., and Scheres, S.H. (2014). Cryo-EM structure of the Plasmodium falciparum 80S ribosome bound to the anti-protozoan drug emetine. Elife 3. 10.7554/eLife.03080.

Wu, B., Eliscovich, C., Yoon, Y.J., and Singer, R.H. (2016). Translation dynamics of single mRNAs in live cells and neurons. Science 352, 1430–1435. doi:10.1126/science.aaf1084.

Wu, C.C., Peterson, A., Zinshteyn, B., Regot, S., and Green, R. (2020). Ribosome Collisions Trigger General Stress Responses to Regulate Cell Fate. Cell 182, 404–416 e414. 10.1016/j.cell.2020.06.006.

Yan, L.L., Simms, C.L., McLoughlin, F., Vierstra, R.D., and Zaher, H.S. (2019). Oxidation and alkylation stresses activate ribosome-quality control. Nat Commun 10, 5611. 10.1038/s41467-019-13579-3.

Yan, X., Hoek, T.A., Vale, R.D., and Tanenbaum, M.E. (2016). Dynamics of Translation of Single mRNA Molecules In Vivo. Cell 165, 976–989. 10.1016/j.cell.2016.04.034.

Yanagitani, K., Kimata, Y., Kadokura, H., and Kohno, K. (2011). Translational pausing ensures membrane targeting and cytoplasmic splicing of XBP1u mRNA. Science 331, 586–589. 10.1126/science.1197142.

Yip, M.C.J., and Shao, S. (2021). Detecting and Rescuing Stalled Ribosomes. Trends Biochem Sci 46, 731–743. 10.1016/j.tibs.2021.03.008.

Zhao, T., Chen, Y.M., Li, Y., Wang, J., Chen, S., Gao, N., and Qian, W. (2021). Disome-seq reveals widespread ribosome collisions that promote cotranslational protein folding. Genome Biol 22, 16. 10.1186/s13059-020-02256-0.

## References

Boersma, S., Khuperkar, D., Verhagen, B.M.P., Sonneveld, S., Grimm, J.B., Lavis, L.D., and Tanenbaum, M.E. (2019). Multi-Color Single-Molecule Imaging Uncovers Extensive Heterogeneity in mRNA Decoding. Cell 178, 458–472 e419. 10.1016/j.cell.2019.05.001.

